# Cortical microtubule remodelling during strigolactone- and light-mediated growth inhibition of Arabidopsis hypocotyls

**DOI:** 10.1101/2020.12.07.414524

**Authors:** Yuliya A. Krasylenko, George Komis, Sonya Hlynska, Tereza Vavrdová, Miroslav Ovečka, Tomáš Pospíšil, Jozef Šamaj

## Abstract

Strigolactones are phytohormones involved in shoot branching and hypocotyl elongation. The latter phenomenon was addressed herein by the exogenous application of a synthetic strigolactone GR24 and an inhibitor of strigolactone biosynthesis TIS108 on hypocotyls of wild type Arabidopsis and a strigolactone signalling mutant *max2-1 (more axillary growth 2-1)*. Owing to the interdependence between light and strigolactone signalling, the present work was extended to seedling cultivation under a standard light/dark regime, or under continuous darkness. Given the essential role of the cortical microtubules in cell elongation, their organization and dynamics were characterized under the conditions of altered strigolactone signalling using fluorescence microscopy methods with different spatiotemporal capacities such as confocal laser scanning microscopy and structured illumination microscopy. It was found that the strigolactone-dependent inhibition of hypocotyl elongation correlated with changes in cortical microtubule organization and dynamics, visualized in living wild type and *max2-1* seedlings stably expressing genetically-encoded fluorescent molecular markers for microtubules. Quantitative analysis of microscopic datasets revealed that chemical and/or genetic manipulation of strigolactone signalling affected microtubule remodelling, especially under light conditions. The application of GR24 and TIS108 in dark conditions partially alleviated cytoskeletal rearrangement, suggesting a new mechanistic connection between the cytoskeletal behaviour and the light-dependence of strigolactone signalling.

**Highlight:** Strigolactones regulate organization and dynamics of cortical microtubules in hypocotyl cells, which contributes to the light-mediated inhibition of hypocotyl growth in Arabidopsis seedlings.

## Introduction

Strigolactones (SL), the carotenoid-derived plant hormones and rhizosphere signaling molecules, were discovered in exogenous allelochemical responses as germination stimulants of Orobanchaceae root parasitic weed (*Striga, Orobanche, Phelipanche*, and *Alectra* spp.) (Cook *et al*., 1966; Gomez-Roldan *et al*., 2008; Koltai, 2014). SL induce hyphal branching of arbuscular mycorrhizal fungi (Akiyama *et al*., 2005), promote nodulation in the legume-rhizobium symbiosis (Soto *et al*., 2009), and enhance plant resistance to drought, salt and osmotic stresses, and to low soil phosphate and nitrate content (Yoneyama *et al*., 2007; Foo *et al*., 2013; Ha *et al*., 2014). The physiological effects of SL on the aboveground plant part include the regression of plant height and hypocotyl length (Stirnberg *et al*., 2002; de Saint Germain *et al*., 2013), regulation of shoot branching by modulating auxin transport (Kapulnik *et al*., 2011; Shinohara *et al*., 2013), increased expansion of cotyledons in etiolated Arabidopsis seedlings (Stirnberg *et al*., 2002; Tsuchiya *et al*., 2010), suppression of the preformed axillary bud outgrowths (Gomez-Roldan *et al*., 2008; Umehara *et al*., 2008; Domagalska and Leyser, 2011), rescue of the dark-induced elongation of rice mesocotyls (Hu *et al*., 2010), promotion of the secondary growth and cell divisions in cambium, and stimulation of leaf senescence (Agusti *et al*., 2011; Koren *et al*., 2013; Koltai, 2014). Furthermore, synthetic SL GR24 acts synergistically with auxins and it is used in potato tuber formation, the outgrowth of the axillary stolon buds, and above-ground shoot branching (Roumeliotis *et al*., 2012). Not only parasitic plant seeds undergo enhanced maturation in SL presence, but also Arabidopsis seeds germinate faster (Tsuchiya *et al*., 2010).

SL are perceived by the α/β-hydrolase receptor DWARF14 (Seto *et al*., 2019) via a specific receptor system, while subsequent signalling requires the SKP1-CULLIN-F-BOX (SCF) complex and proteasome-mediated degradation of target proteins (reviewed by Kumar *et al*., 2015a; Yoneyama *et al*., 2020). At physiological and molecular levels, SL have been suggested to affect auxin efflux (Koltai, 2014) through PIN1 efflux carrier in the root (Ruyter-Spira *et al*., 2011) but also to dampen auxin transport in the shoot (Domagalska and Leyser, 2011).

Plant cytoskeleton is involved in many processes regulated by SL, e.g. the switch from the cell division to expansion (Ruan and Wasteneys, 2014), the cell elongation and differentiation (Ivakov and Persson, 2013; Ambrose and Wasteneys, 2014; Sampathkumar *et al*., 2014), as well as in plant responses to salt (Shoji *et al*., 2006) and osmotic stresses (Komis *et al*., 2002; Wang *et al*., 2010).

It is presumed that SL, as a new class of phytohormones, indirectly regulate cytoskeleton organization together with well-studied phytohormones such as auxins, cytokinins, giberellins and abscisic acid. Through the indirect regulation of microtubules (MT) and actin filaments, SL should orchestrate morphogenesis of both above- and underground plant parts (Blume *et al*., 2017). Previously, it was reported that SL affect architecture and dynamics of actin filaments in Arabidopsis root cells (Pandya-Kumar *et al*., 2014). GR24 reduces F-actin filament bundling in a MORE AXILLARY GROWTH2 (MAX2)-dependent manner and, at the same time, enhances actin dynamics, affects endosome trafficking and PIN2 localization in the plasma membrane (Pandya-Kumar *et al*., 2014). Moreover, plant response to low phosphorus conditions involves MAX2-dependent reduction of PIN2 and endosome trafficking, plasma membrane polarization, and increased actin filament bundling in epidermal root cells (Kumar *et al*., 2015b). Concerning MT, the other fundamental cytoskeletal component, SL analogues MEB55 and ST362 were found to compromise the integrity of the MT network in animal cells (Mayzlish-Gati *et al*., 2015). Moreover, SL analogues also affect the MT regulation by activating apoptotic p38 (mitogen-activated protein kinase) cascade and inhibiting cyclin B expression (Pollock *et al*., 2014).

However, there are no reports on the effects of SL on plant MT so far. This study shows effects of exogenously applied synthetic SL GR24 and inhibitor of SL biosynthesis TIS108 on the organization and dynamics of cortical MT in epidermal cells of light-exposed and etiolated hypocotyls of wild type plants and SL-insensitive Arabidopsis mutant *max2-1*. Our results suggest that strigolactones affect mainly MT cytoskeleton under light conditions and this effect can be alleviated by darkness. From the methodological point of view, we have used combination of confocal laser scanning confocal microscopy (CLSM) and super-resolution structured illumination microscopy (SIM) to obtain these results.

## Materials and methods

### Plant material and growth conditions

Wild-type *Arabidopsis thaliana* plants ecotype Columbia-0 (Col-0), and the SL-insensitive *A. thaliana max2-1* mutant (EMS mutant in Col-0-background; Stirnberg *et al*., 2002) kindly provided by Prof. H. Koltai, were used in this study. MT dynamics were recorded in seedlings stably expressing a *35S::GFP-MBD* (MT-binding domain of mammalian non-neuronal MICROTUBULE ASSOCIATED PROTEIN4), or a *35S::TUA6-GFP* construct (α-TUBULIN 6) (Marc *et al*., 1998; Shaw *et al*., 2003). Mutant plants of *max2-1* were crossed with lines carrying *35S::GFP-MBD* or *35S::TUA6-GFP* constructs. For microscopy studies the F3 generation of the progenies was used. Homozygous *max2-1* seedlings expressing the above MT markers were selected according to fluorescence detection under epifluorescence microscope.

Prior to germination, seeds were sterilized in 1% v/v sodium hypochlorite solution supplemented with 0.1% v/v Tween-20 for 10 min, short-spin vortexed, immersed to 70% v/v ethanol for 5 s, thoroughly rinsed 5 times by MilliQ water and placed to 0.6% w/v agarose-solidified ½ Murashige and Skoog medium (½ MS; Duchefa, the Netherlands) with 10% w/v sucrose with or without exogenous synthetic SL and/or inhibitors of endogenous SL biosynthesis.

### Chemical treatment

Unless stated otherwise, all common chemicals were from Sigma-Aldrich (the USA) and were of analytical grade. A synthetic specific SL *cis-*GR24 consisting only of D14-perceived GR24+ was synthesized according to Zwanenburg *et al*., 2013 was dissolved *ex tempore* in pure anhydrous acetone to prepare a 10 mM stock solution from which working concentrations of 3 and 25 μM were prepared. Four-day-old seedlings were taken from 0.6% w/v agarose-solidified media, treated with GR24 and prepared for microscopy. A triazole-type SL biosynthesis inhibitor designated as TIS108 (Chiralix, the Netherlands) was dissolved in pure anhydrous acetone prior to use to obtain 10 mM stock solution further diluted to 3 μM final concentration and added to agarose-solidified ½ MS medium or used for short-time treatment in liquid MS. Petri dishes with seeds were stored at 4°C for 1 day to synchronize germination and then germinated at a vertical position in Phytotron at 22°C under long-day conditions (16 h light/8 h darkness, photosynthetic photon flux (PPF): 150 µmol m^−2^ s^−1^) for 4 or 7 days prior to imaging. For the etiolation experiment, Petri dishes were wrapped in aluminum foil after seeding, stratified at 4^°^C for 2–4 days, and germinated as such under the same environmental conditions.

### Hypocotyl growth analysis

Petri dishes with 3-7 day old seedlings were placed in scanner (Image Scanner III, Seiko Epson, Japan) and scanned at transmitted light mode in order to document and subsequently quantify hypocotyl length. For hypocotyl width measurements, seedlings were documented with differential interference contrast of a widefield microscope (Axio Imager M2, Carl Zeiss, Germany) equipped with a polarizer and a Wollaston prism at three distinct parts of the hypocotyl: the upper part – situated right beneath the cotyledon petiole; the middle part – at the mid-plane of hypocotyl, and the lower part– at the border with the primary root.

For the detailed morphological studies 4- and 7-days old seedlings were captured using Axio Zoom.V16 Stereo Zoom system (Carl Zeiss, Germany) in bright field illumination (objective lenses PlanApo Z 1.5x, FWD = 30mm). The measurements were done using the default Measure application of ImageJ (Schneider et al. 2012) by tracking hypocotyls with the segmented line tool after appropriate scale calibration using the Set Scale tool of the Analyze menu.

### Microscopy

For live imaging of MTs four different Zeiss microscopy platforms (Zeiss Microscopy, Germany) were used (Komis et al., 2014; 2015). For deciphering MT organization, GFP-MBD or TUA6-GFP molecular markers were visualized by means of CLSM with the LSM710 system (Carl Zeiss, Germany) equipped with a 63× Plan-Apochromat oil-immersion objective (1.4 NA) under excitation 488/543 nm, emission 510/540 nm. Laser excitation intensity did not exceed 2% of the laser intensity range available. Range of the Z-stack was always set up to 0.61 µm. GFP-labelled MT were imaged using excitation laser line 488 nm and emission spectrum 493-630 nm for GFP fluorescence detection and with excitation laser line 405 nm and emission spectrum 410-495 nm for DAPI fluorescence detection.

Microscopy platform enabling SIM (ELYRA PS.1, Carl Zeiss, Germany) with 63× Plan-Apochromat oil-immersion objective was used for the time-lapse observations of MT dynamics. 4-days old seedlings were mounted between a microscope slide and a coverslip in 30 μL of liquid MS medium spaced by double-sided sticky tape, narrow Parafilm stripes and extra sealed using liquid petroleum jelly (nail polish) to form a chamber prior to imaging for sample stabilization. This prevented dislocation of the plantlets during liquid exchange and allowed the observation of the same area during 2 h. Seedlings were grown at solidified GR24/TIS108- containing media for 4 d.

All preparations with the etiolated seedlings were done quickly in dark room using dim red or green light to prevent disturbances of MT by visible light.

### Post-acquisition image processing

Raw SIM images were processed automatically by the respective add-on of the licensed Zen software (Black version; Carl Zeiss, Germany) coupled to the Elyra PS.1, according to standards thoroughly described before (Komis et al., 2014; 2015).

Kymographs of MT time series recordings were generated using the Kymograph add-on of the licensed Zen software (Blue version; Carl Zeiss, Germany), using the arrow tool to delineate individual or bundled MT of interest.

### Quantitative analysis of microtubule organization

MT organization was quantitatively addressed by assessing the extend of MT bundling as the skewness of fluorescence distribution of GFP-MBD expressing cells. Skewness was extrapolated from values provided by the Histogram add-on of licensed Zen software (Blue version). Additionally, cortical MT ordering was quantitatively assessed by measuring MT organization anisotropy. This anisotropy was demonstrated in full frames of CSLM images of hypocotyl cells expressing GFP-MBD, which were analyzed with Cytospectre freeware (Kartasalo *et al*., 2015) to extrapolate their angular distribution in the cortical cytoplasm. Quantitatively, the ordering of cortical MT was measured through the FibrilTool macro as described previously (Boudaoud *et al*., 2014). Briefly, the FibrilTool macro was applied on regions of interests drawn using the Polygon tool of Image J delineating the circumference of fully visible cells. Care was taken, to avoid cell edges, where frequently the signal is saturated. Theoretically the numerical result ranges between 0 (complete isotropy) to 1 (perfect anisotropy).

### Quantitative analysis of microtubule dynamics

Kymographs from recordings of dynamic MT were used to extrapolate the following parameters of MT dynamics: growth and shrinkage rates, catastrophe and rescue frequencies. Kymograph analysis was done manually using the Image J angle measure tool after size calibration of kymographs. Angles were acquired in degrees and converted to radians in MS Excel (Microsoft, the USA) prior to calculations of tangential values. Briefly, the equations used were as follows:

For growth rate, the equation was: *G* = tan *φ* × *pixel size* × *fps*; where tan_*φ*_ is the tangential of the growth slope, pixel size is in µm and fps is the frame rate of the acquisition (frames×sec^-1^). The final output is converted to µm×min^-1^ by multiplying the original value with 60 sec×min^-1^.

For shrinkage rate, the equation was: *S* = tan *θ* × *pixel size* × *fps*; tan_*θ*_ is the tangential of the shrinkage slope, pixel size is in µm and fps is the frame rate of the acquisition (frames×sec^-1^). The final output is converted to µm×min^-1^ by multiplying the original value with 60 sec×min^-1^.

For catastrophe frequency the following equation was applied:

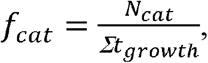

where f_*cat*_ is the catastrophe frequency, N_*cat*_ is the total number of catastrophe events and ∑t_*growth*_ is the total time spent in growth, regarding all the growth events taken into account.

For rescue frequency the following equation was applied:

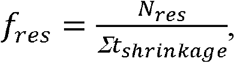

where f_*res*_ is the rescue frequency, N_*res*_ is the total number of rescue events and ∑t_*growth*_ is the total time spent in shrinkage, regarding all the shrinkage events taken into account.

### Statistics

Statistical analysis of all datasets was performed in the software STATISTICA (version 13.4.0.14; Statsoft, the USA). All datasets were first subjected to Shapiro-Wilk W test and Levene’s tests to test the normality and homogeneity. Frequently, the datasets failed to pass these tests. On several representative datasets, following tests were calculated: (i) one-way ANOVA, (ii) Welch’s ANOVA, (iii) Tukey’s post hoc test corrected for unequal sample size, (iv) Scheffé’s post hoc test, (v) Kruskal-Wallis test. Based on results of these preliminary analysis, and in agreement with previous reports (Liu, 2015), Welch’s ANOVA followed by Scheffé’s test was used as it exhibited higher stringency compared to other tests. Statistical significance was determined based on the calculated p-values, were for Welch’s ANOVA the probability level was 0.05 and for Scheffé’s test the probability level was 0.01. For comparing two different experimental conditions (light-dark and inhibitor treatment), two-way ANOVA followed by Scheffé’s test was used. In this case, the probability level set for Scheffé’s test was 0.001.

## Results

### Strigolactone treatment affects hypocotyl growth in Arabidopsis

Col-0 and *max2-1* mutant seedlings were germinated and cultivated on medium containing different concentrations of GR24 or TIS108, either in the standard light/dark regime of the phytotron or in the dark. Under the light/dark regime, GR24 used at two different concentrations (3 and 25 µM), stalled hypocotyl elongation and induced mild radial swelling as compared to mock-treated Col-0 (Fig. 1A,I cf. Fig. 1B,C,J,K). Growth inhibition of the hypocotyl was also evident after treatment with the inhibitor TIS108 tested in two concentrations (Fig. 1D,L). In quantitative terms, the hypocotyl length of mock treated Col-0 seedlings was 2.05±0.145 mm (mean±SD; Fig. 1Q; N=75; Supplementary Table S1). After treatment with 3 µM GR24, the hypocotyl length was significantly reduced to 1.28±0.158 mm (mean±SD; Fig. 1Q; p=0.0000; N=60), while after treatment with 25 µM the hypocotyl length comprised 1.307±0.178 (mean±SD; Fig. 1Q; N=56), which was significantly different compared to mock-treated Col-0, but not to the effect of 3 µM GR24 (p=0.0000 and p=0.9992, respectively). In turn, TIS108 treatment resulted in the most severe hypocotyl growth inhibition measured to 0.783±0.160 mm (mean±SD; Fig. 1Q; N=61), being significantly different from all other conditions tested (p=0.0000 as compared to treatment with 3 and 25 µM GR24). Hypocotyl width was only slightly affected by any of the treatments used herein. Briefly the width of mock-treated Col-0 hypocotyls was 0.309±0.04 mm (Fig. 1R; N=58; Supplementary Table S2), 0.325±0.07 mm after treatment with 3 µM GR24 (Fig. 1R; N=89), 0.290±0.08 mm after treatment with 25 µM GR24 (Fig. 1R; N=90) and 0.323±0.05 mm after treatment with 3 µM TIS108.

**Fig. 1.**
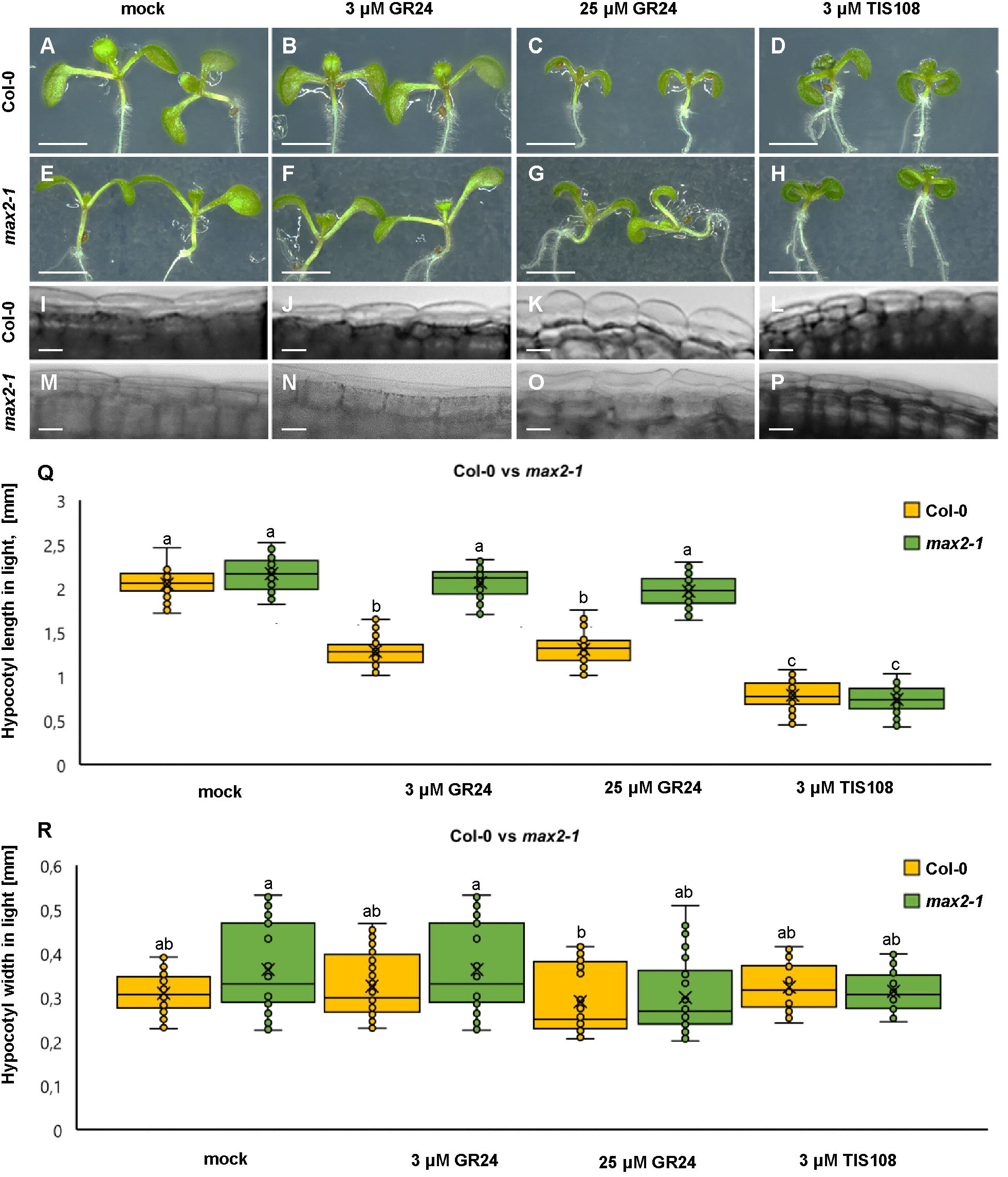
Hypocotyl development of light-grown seedlings of Arabidopsis Col-0 or the *max2-1* mutant in the presence or absence of GR24 synthetic strigolactone (3 and 25 µM) or the biosynthetic inhibitor of strigolactone production TIS108 (3 µM). (A–D). Overview of hypocotyl of Col-0 seedlings treated with solvent alone (mock; A); 3 µM GR24 (B); 25 µM GR24 (C); 3 µM TIS108 (D). (E–H) Similar overview of hypocotyls of light-grown *max2-1* mutant seedlings in the presence of solvent alone (mock; E), 3 µM GR24 (F); 25 µM GR24 (G); 3 µM TIS108 (H). (I–L) Magnified views of hypocotyls of Col-0 treated with solvent alone (mock; I); 3 µM GR24 (J); 25 µM GR24 (K); 3 µM TIS108 (L) showing mild cell swelling in all treatments (J–L) compared to control (I). (M–P) Similar comparison of *max2-1* hypocotyl epidermal cells treated with solvent alone (mock; M); 3 µM GR24 (N); 25 µM GR24 (O); 3 µM TIS108 (P). (Q) Quantitative assessment of Col-0 and *max2-1* hypocotyl length comparing pairwise mock treatment and treatments with 3 µM GR24, 25 µM GR24, and 3 µM TIS108 (N≥59; two-way ANOVA was followed with Scheffé’s test, statistical comparison is shown within groups sharing the same genotype; letters in the graph are shared by groups without statistically significant differences at the 0.001 probability level; results are in Table S1). (R) Quantitative assessment of Col-0 and *max2-1* hypocotyl width comparing pairwise mock treatment and treatments with 3 µM GR24, 25 µM GR24 and 3 µM TIS108 (N≥27; two-way ANOVA was followed with Scheffé’s test, statistical comparison is shown within groups sharing the same genotype; letters in the graph are shared by groups without statistically significant differences at the 0.001 probability level; results are in Table S2). In all box plots, average is presented by ×, median by the middle line, 1st quartile by the bottom line, 3^rd^ quartile by the top line; the whiskers lie within the 1.5× interquartile range (defined from the 1^st^ to the 3^rd^ quartile) while outliers are omitted. Scale bars: 5 mm (A–H); 5 µm (I–P).

Subsequently we characterized hypocotyl growth in light-exposed *max2-1* mutants. In such mock-treated mutants (Fig. 1E) as well as after the treatment with both 3 (Fig. 1F) and 25 µM GR24 (Fig. 1G), the hypocotyl length was comparable to mock-treated Col-0 seedlings. The hypocotyl length of *max2-1* mutant seedlings was only responsive to treatment with 3 µM TIS108 (Fig. 1H). Meanwhile, hypocotyl width does not show any noticeable changes at any treatment used (Fig. 1M–P). In quantitative terms, the hypocotyl length of mock-treated *max2-1* mutants was 2.16±0.19 mm (mean±SD; N=78), 2.06±0.17 mm after 3 µM GR24 (mean±SD; N=69), 1.98±0.17 mm after 25 µM GR24 (mean±SD; N=80), and 0.74±0.16 mm after 3 µM TIS108 (mean±SD; N=59) treatments. The hypocotyl length of *max2-1* mutants was significantly different at all GR24 and TIS108 treatments as compared to the treated Col-0 seedlings (Fig. 1Q; p=0.185 after mock treatment, p=0.9999 after 3 µM GR24, p=0.2333 after 25 µM GR24, and p=0.0000 after 3 µM TIS108 treatments). Within the *max2-1* population, GR24 treatments did not affect hypocotyl length (Fig. 1Q) and only treatment with 3 µM TIS108 brought about its significant shortening (Fig. 1Q). In terms of hypocotyl width, no changes were discerned (Fig. 1R).

### Strigolactone effects are modulated in dark-grown seedlings

There have been reports of a synergy between exogenous application of SL and the illumination conditions during seedling growth. Therefore, the experimental regime of the treatment of Col-0 and *max2-1* seedlings with two different concentrations of GR24 (3 and 25 µM) and with 3 µM TIS108 grown under persistent darkness was tested.

As expected, hypocotyl length elongation of etiolated seedlings exceeds that of light-grown seedlings (Fig. 2A). By combining visual documentation and quantitative analysis, it became evident that etiolated seedlings were also responsive to treatments with 3 (Fig. 2B,I) and 25 µM GR24 (Fig. 2C,I), but were the most sensitive to 3 µM TIS108 (Fig. 2D,I). The length of mock-treated etiolated Col-0 seedlings was 15.45±1.77 mm (mean±SE; N=78), 14.09±1.22 mm after treatment with 3 µM GR24 (mean±SE; N=75), 11.23±1.54 mm after 25 µM GR24 (mean±SE; N=77), and 2.55±0.72 mm after 3 µM TIS108 (N=13). All the treatments caused significant reduction of the etiolated hypocotyl length as compared to the mock treatment (Fig. 2I; p=0.0000 for 3 µM GR24; p=0.0000 for 25 µM GR24; p=0.0000 for 3 µM TIS108; Supplementary Table S3).

**Fig. 2.**
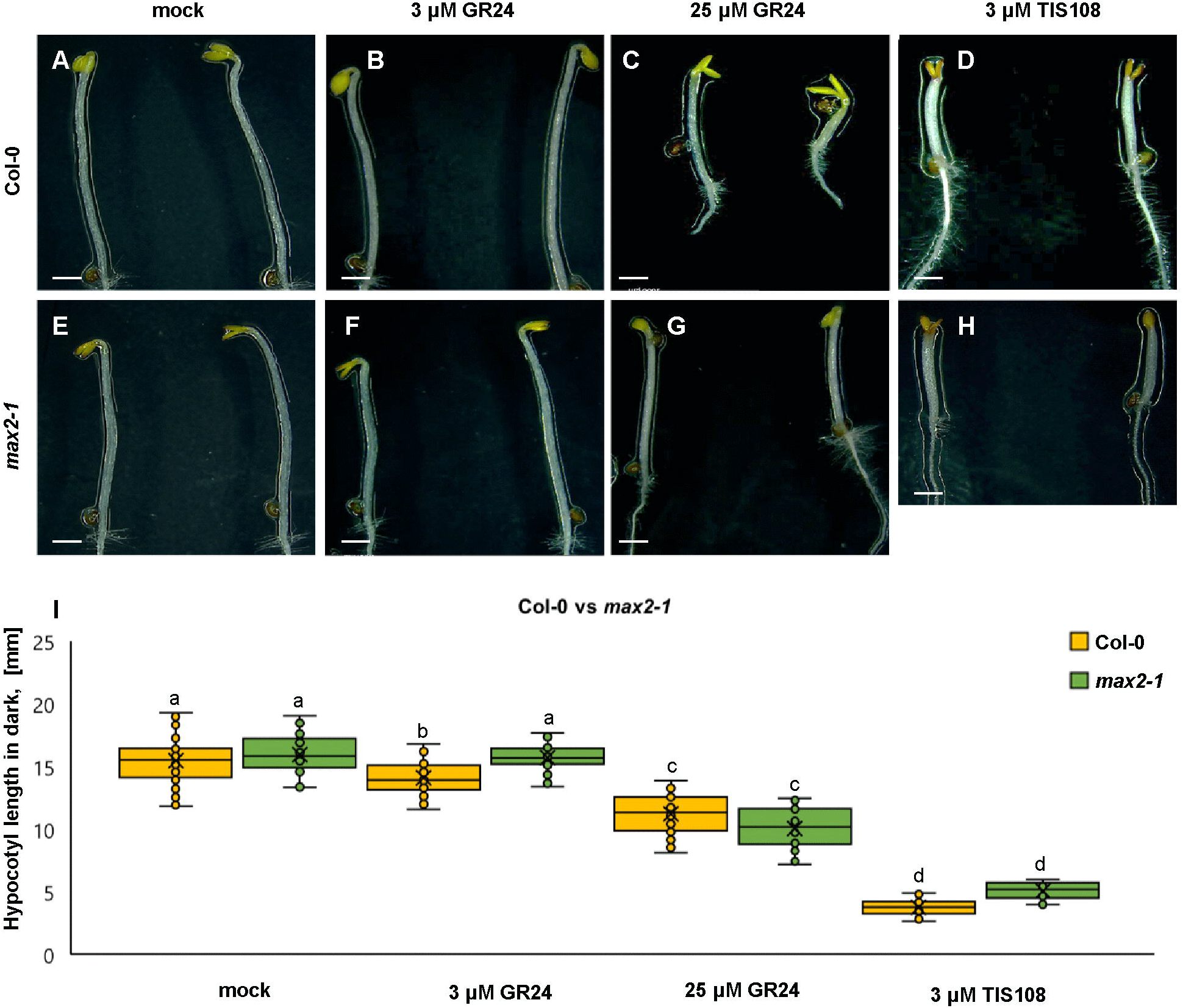
Hypocotyl development of dark-grown seedlings of Arabidopsis Col-0 or the *max2-1* mutant in the presence or absence of GR24 synthetic strigolactone (3 µM and 25 µM) or the biosynthetic inhibitor of strigolactone production TIS108 (3 µM). (A–E). Overview of etiolated hypocotyl of Col-0 seedlings treated with solvent alone (mock; A); 3 µM GR24 (B); 25 µM GR24 (C); 3 µM TIS108 (D). (E–H) Similar overview of hypocotyls of etiolated *max2-1* mutant seedlings in the presence of solvent alone (mock; E); 3 µM GR24 (F); 25 µM GR24 (G); 3 µM of TIS108 (H). (I) Quantitative assessment of etiolated Col-0 and *max2-1* hypocotyl length comparing pairwise mock treatment and treatments with 3 µM GR24, 25 µM GR24, and 3 µM TIS108 (N≥22; two-way ANOVA was followed with Scheffé’s test, statistical comparison is shown within groups sharing the same genotype; letters in the graph are shared by groups without statistically significant differences at the 0.001 probability level; results are in Table S3). In all box plots, average is presented by ×, median by the middle line, 1^st^ quartile by the bottom line, 3^rd^ quartile by the top line; the whiskers lie within the 1.5^×^ interquartile range (defined from the 1^st^ to the 3^rd^ quartiles) while outliers are omitted. Scale bars: 10 mm (A–H).

The same line of experiments was applied in the case of *max2-1* mutants (Fig. 2E-H), which also proved to be prone to either the application of exogenous SL, or to the metabolic inhibition of SL biosynthesis. Compared to mock treated *max2-1* seedlings (Fig. 2E,I), which exhibited length of 15.95±1.64 mm (mean±SD; N=23), those treated with 3 µM GR24 (Fig. 2F,I) were 15.72±1.12 mm (mean±SD; N=30), those treated with 25 µM GR24 (Fig. 2G,I) were 10.05±1.66 mm (mean±SD; N=22), while those treated with 3 µM TIS108 (Fig. 2H,I) were 3.07±0.66 mm (mean±SD; N=19). The effect of most treatments on hypocotyl length of etiolated *max2-1* seedlings was deemed to be significant by comparison to the mock treatment except for 3 µM GR24 (Fig. 2I; p=0.9998 for 3 µM GR24; and p=0.0000 for both 25 µM GR24 and 3 µM TIS108).

Such preliminary deductions make the assessment of whether light plays a crucial role on the modulation of SL effects. For this reason, we examined the extent of its effect on the percentage of hypocotyl reduction of either light- or dark-grown Col-0 or *max2-1* mutant seedlings by comparison to the mock treatment. Thus, under the light exposure combined with 3 µM GR24 treatment the reduced hypocotyl length was observed, comprising 63.33±9.68% in Col-0 (Supplementary Fig. 1A) and 93.01±12.48% in etiolated seedlings (Supplementary Fig. 1B) as compared to mock-treated seedlings. Similarly, the treatment with 25 µM GR24 caused the reduction of hypocotyl length up to 64.35%±9.63% of mock-treated seedlings under light exposure, but this reduction was less pronounced in dark (cf. 74.00%±13.69% of mock treated seedlings). TIS108 treatment caused hypocotyl length reduction in light up to 38.55%±8.17% of mock-treated seedlings, and even more pronounced reduction in the dark, since the treated etiolated hypocotyls were 23.90%±4.61% of the mock-treated counterparts.

Regarding the light-exposed *max2-1* mutant, hypocotyl length of seedlings treated with 3 µM GR24 (Supplementary Fig. 1C) was 96.65%±11.09% of mock-treated seedlings, of those treated with 25 µM GR24 – 91.88%±10.55%, and of those treated with 3 µM TIS108 – 34.72%±8.02%, respectively. Moreover, hypocotyl length of etiolated *max2-1* seedlings (Supplementary Fig. 1D) after the treatment with 3 µM GR24 was 97.38%±6.92% of mock-treated seedlings, of those treated with 25 µM GR24 – 63.18%±6.39%, and of those treated with 3 µM TIS108 – 19.74%±3.19%, respectively.

These results revealed the extent of GR24 and TIS108 effects on hypocotyl elongation, showing that etiolated Col-0 seedlings are less responsive to GR24 as compared to light-grown ones, while at the same time they were more sensitive to TIS108 treatment. In addition, *max2-1* mutants were equally unresponsive to 3 µM GR24, however, both 25 µM GR24 and 3 µM TIS108 caused much stronger inhibitory effect in etiolated seedlings.

### Strigolactone affects microtubule organization in light-dependent manner

Inducible growth alterations following extrinsic stimulation as, e.g., with hormonal treatments, has been repeatedly shown to be preceded and supported by conditional rearrangements of cortical MT, which tend to keep the predominant orientation (e.g., Lindeboom *et al*., 2013; True and Shaw, 2020). Such conditions favouring the parallel arrangement of cortical MT can be documented by showing the patterns of their angular distribution and quantified by measuring the degree of anisotropy within the cortical array.

In light-grown, mock-treated Col-0 seedlings expressing a GFP-MBD MT marker, cortical MT exhibit a more or less random distribution (Fig. 3A,E) with the tendency of more biased reorganization after treatment with 3 (Fig. 3B,F) and 25 µM GR24 (Fig. 3C,G), and 3 µM TIS108 (Fig. 3D,H). By contrast, *max2-1* mutants expressing the same MT marker appeared to have more organized cortical MT as compared to Col-0 either after mock treatment (Fig. 3I,M) and treatments with 3 (Fig. 3J,N) and 25 µM GR24 (Fig. 3K,O), and 3 µM TIS108 (Fig. 3L,P).

**Fig. 3.**
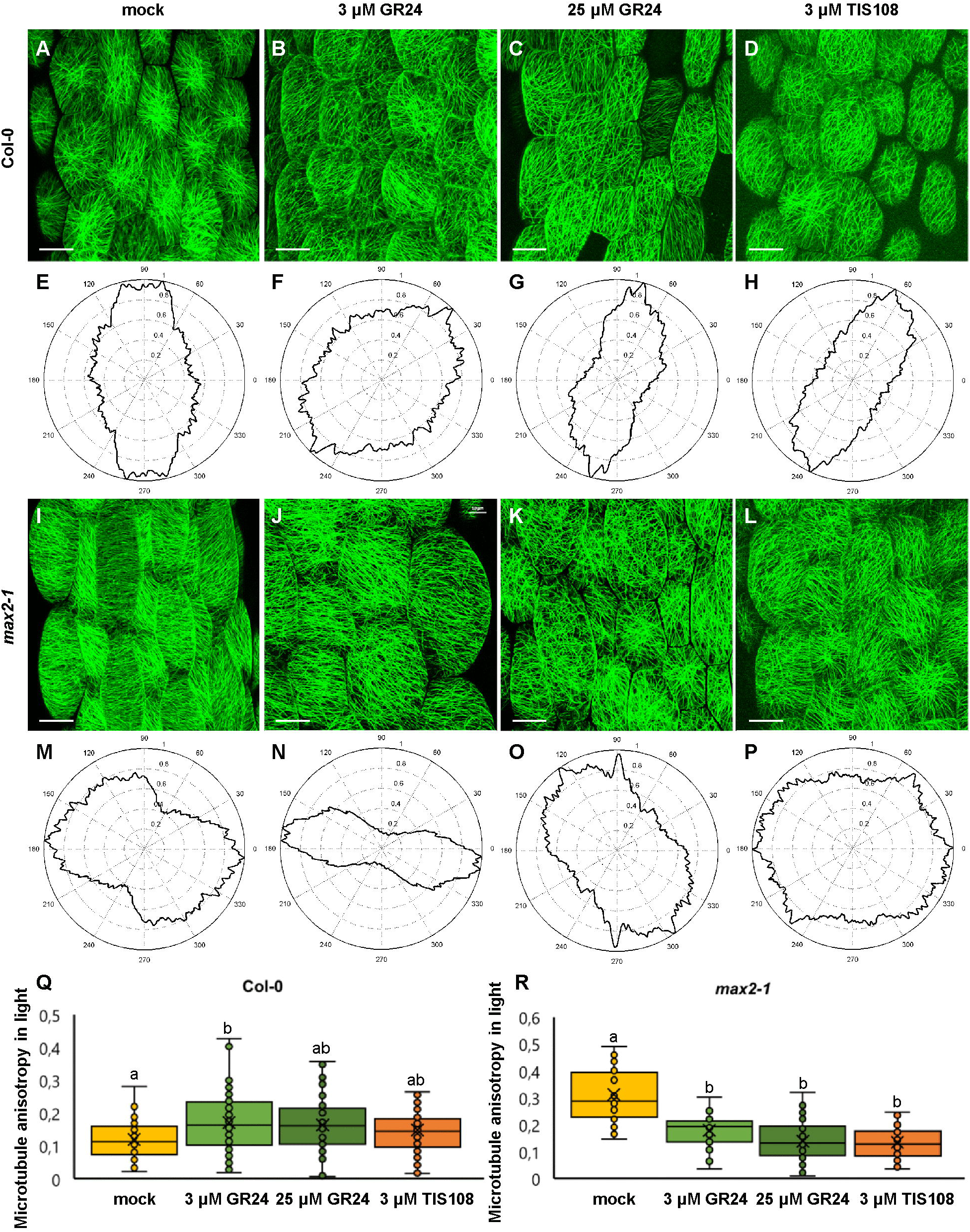
Assessment of microtubule organization in epidermal hypocotyl cells of light-grown seedlings of Arabidopsis Col-0 or the *max2-1* mutant in the presence or absence of GR24 synthetic strigolactone (3 and 25 µM) or the biosynthetic inhibitor of strigolactone production TIS108 (3 µM). (A–D). Overview of hypocotyl of Col-0 seedlings treated with solvent alone (mock; A), 3 µM GR24 (B); 25 µM GR24 (C); 3 µM TIS108 (D). (E–H) Cytospectre graphs of cortical microtubule distribution where (E) corresponds to (A), (F) to (B), (G) to (C), and (H) to (D). (I–L) Overview of hypocotyl of *max2-1* seedlings treated with solvent alone (mock; I); 3 µM GR24 (J); 25 µM GR24 (K); 3 µM TIS108 (L). (M–P) Cytospectre graphs of cortical microtubule distribution where (M) corresponds to (I), (N) to (J), (O) to (K), and (P) to (L). (Q) Quantitative assessment of anisotropy of cortical microtubule organization in light- grown Col-0 after mock treatment and treatments with 3 µM GR24; 25 µM GR24 and 3 µM TIS108 (N≥54; Welch’s ANOVA was followed with Scheffé’s test, statistical comparison is shown within groups sharing the same genotype; letters in the graph are shared by groups without statistically significant differences at the 0.01 probability level; results are in Table S4). (R) Quantitative assessment of anisotropy of cortical microtubule organization in light-grown *max2-1* after mock treatment and treatments with 3 µM GR24, 25 µM GR24, and 3 µM TIS108 (N≥31; Welch’s ANOVA was followed with Scheffé’s test, statistical comparison is shown within groups sharing the same genotype; letters in the graph are shared by groups without statistically significant differences at the 0.01 probability level; results are in Table S5). In all box plots, average is presented by ×, median by the middle line, 1st quartile by the bottom line, 3^rd^ quartile by the top line; the whiskers lie within the 1.5^×^ interquartile range (defined from the 1^st^ to the 3^rd^ quartile) while outliers are omitted. Scale bars: 20 µm.

The qualitative observations mentioned above were quantitatively corroborated by measuring changes in the anisotropy of MT organization. In mock-treated Col-0 anisotropy was 0.12±0.06 (Fig. 3Q; mean±SD; N=54), after treatment with 3 µM GR24 it became 0.17±0.08 (Fig. 3Q; mean±SD; N=71), and 0.016±0.08 (Fig. 3Q; mean±SD; N=54) and 0.15±0.07 (Fig. 3Q; mean±SD; N=75) after 25 µM GR24 and 3 µM TIS108 treatments, respectively. In light-grown Col-0 seedlings all treatments significantly promoted anisotropy within the cortical MT array as compared to the mock treatment (Fig. 3Q; p=0.005 for 3 µM GR24; p=0.0375 for 25 µM GR24; p=0.2381 for 3 µM TIS108).

Oppositely, light-grown mock-treated *max2-1* seedlings exhibited more biased arrays as compared to mock-treated Col-0 seedlings (e.g., Fig. 3I,M), but all treatments (Fig. 3J-P) caused anisotropy reduction to values comparable to those of Col-0, again at significant values (Fig. 3R; p=0.0000 for 3 µM GR24; and p=0.0000 for both 25 µM GR24 and 3 µM TIS108). In this case cortical MT anisotropy was 0.31±0.1 after mock treatment (Fig. 3R; mean±SD; N=35), but was significantly reduced to 0.17±0.07 after treatment with 3 µM GR24 (Fig. 3R; mean±SD; N=31), to 0.14±0.07 after 25 µM of GR25 (Fig. 3R; mean±SD; N=61), and to 0.13±0.06 for 3 µM TIS108 (Fig. 3R; mean±SD; N=79).

In Col-0 etiolated seedlings the degree of cortical MT organization is much more pronounced as compared to light-grown seedlings. In mock-treated seedlings (Fig. 4A,E), cortical MT are largely parallel to each other at variable orientations to the main cell axis, and this pattern seems to be unaffected in seedlings treated with 3 µM GR24 (Fig. 4B,F), 25 µM GR24 (Fig. 4C,G) and 3 µM TIS108 (Fig. 4D,H). Similarly, etiolated seedlings of *max2-1* mutant exhibit highly organized systems of parallel MT, at seemingly the same level of organization comparing to mock-treatment (Fig. 4I,M) or treatments with 3 µM GR24 (Fig. 4J,N), 25 µM GR24 (Fig. 4K,O) and 3 µM TIS108 (Fig. 4L,P). Indeed, this observation was reflected to the level of cortical MT anisotropy, which in Col-0 was statistically similar between all cases (Fig. 4Q; 0.29±0.11, N=27 for mock-treated Col-0; 0.33±0.15, N=42 for Col-0 treated with 3 µM GR24; 0.27±0.12, N=47 for Col-0 treated with 25 µM GR24; and 0.34±0.09 for Col- 0 treated with 3 µM TIS108; mean±SD). In all cases, anisotropy of cortical MT organization was equally high in etiolated seedlings of the *max2-1* mutant with minor variations within this group of treatments (Fig. 4R). Thus, anisotropy values were 0.29±0.08 for mock-treated seedlings (mean±SD; N=29), 0.31±0.14 (mean±SD; N=54) after treatment with 3 µM GR24, 0.31±0.12 (mean±SD; N=51) after 25 µM GR24, and 0.32±0.11 (mean±SD) after 3 µM TIS108 (Fig. 4R). Next, in *max2-1* seedlings only the anisotropy of those treated with 3 µM GR24 was found to be different compared to mock-treated ones (p=0.0280), but not compared to the other treatments, which were similar to the mock. In both cases, it seems that etiolation promotes the biased organization of cortical MT irrespectively of treatments modulating SL activity.

**Fig. 4.**
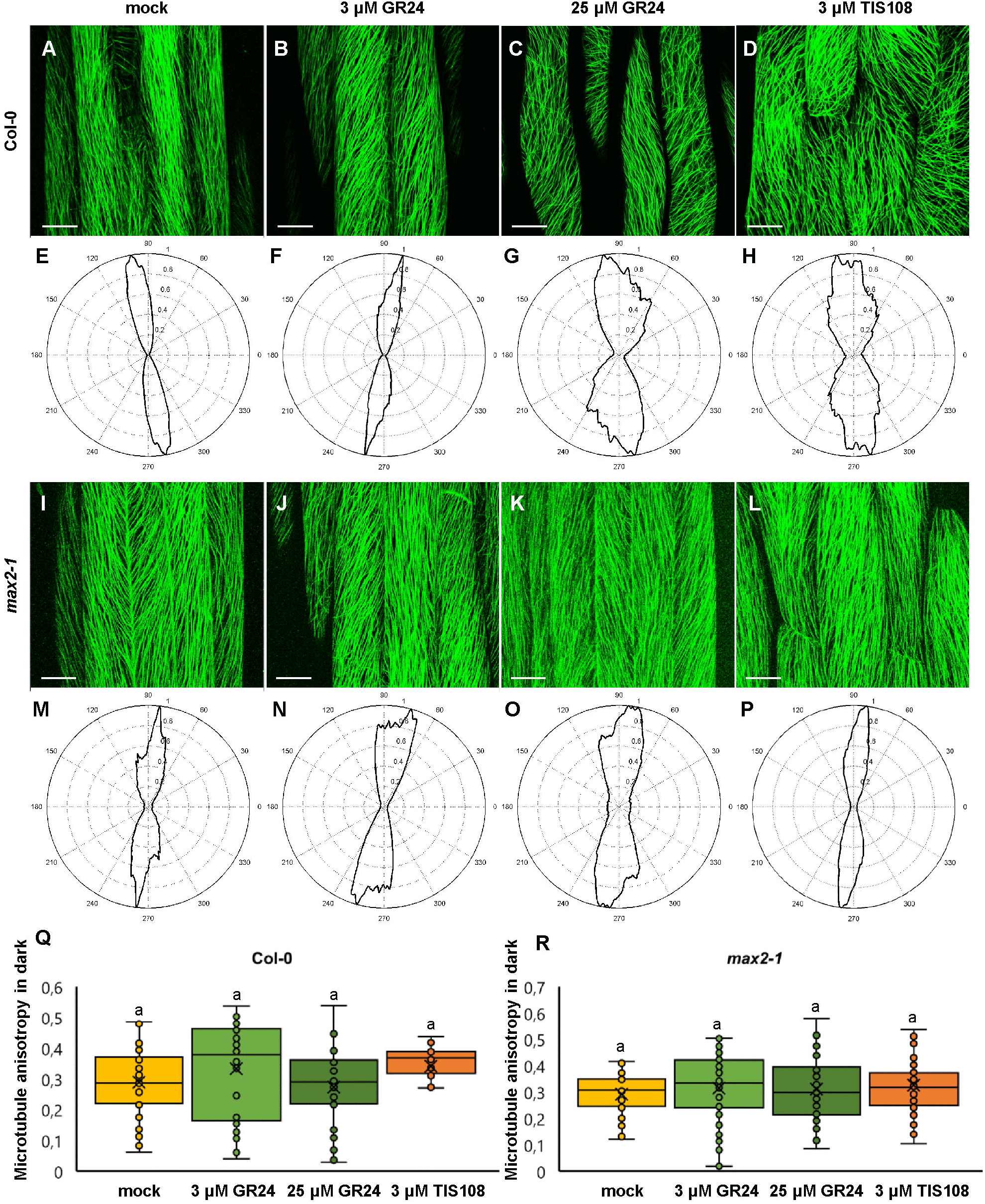
Assessment of microtubule organization in epidermal hypocotyl cells of etiolated seedlings of Arabidopsis Col-0 or the *max2-1* mutant in the presence or absence of GR24 synthetic strigolactone (3 and 25 µM) or the biosynthetic inhibitor of strigolactone production TIS108 (3 µM). (A-D). Overview of hypocotyl of Col-0 seedlings treated with solvent alone (mock; A); 3 µM GR24 (B); 25 µM GR24 (C); 3 µM TIS108 (D). (E–H) Cytospectre graphs of cortical microtubule distribution where (E) corresponds to (A), (F) to (B), (G) to (C), and (H) to (D). (I–L) Overview of hypocotyl of *max2-1* seedlings treated with solvent alone (mock; I); 3 µM GR24 (J); 25 µM GR24 (K); 3 µM TIS108 (L). (M–P) Cytospectre graphs of cortical microtubule distribution where (M) corresponds to (I), (N) to (J), (O) to (K), and (P) to (L). (Q) Quantitative assessment of anisotropy of cortical microtubule organization in light- grown Col-0 after mock treatment and treatments with 3 µM GR24, 25 µM GR24 and 3 µM TIS108 (N≥27; Welch’s ANOVA showed no statistically significant difference within the dataset; F (3, 143)=3.1416, p=0.030; Supplementary Table S4). (R). Quantitative assessment of anisotropy of cortical microtubule organization in light-grown *max2-1* after mock treatment and treatments with 3 µM GR24, 25 µM GR24 and 3 µMTIS108 (N≥29; Welch’s ANOVA was followed with Scheffé’s test, but there was no statistically significant difference at the 0.01 probability level; results are in Table S6). In all box plots, average is presented by ^×^, median by the middle line, 1st quartile by the bottom line, 3^rd^ quartile by the top line; the whiskers lie within the 1.5× interquartile range (defined from the 1^st^ to the 3^rd^ quartile) while outliers are omitted. Scale bars: 20 µm.

### Strigolactone content alterations interfere with microtubule bundling

From the putative mechanisms underlying MT reorganization, bundling is one of the possibilities and it can be related to changes in the distribution of fluorescence intensity frequencies. Uniform labelling results in somewhat normal distribution, while clustered labelling is linked to increasingly skewed distribution, depending on the degree of non-uniformity of the signal. We quantified skewness of fluorescence distribution in hypocotyl cells of either Col-0 or *max2-1* untreated or treated with 3 and 25 µM GR24 or 3 µM TIS108 under light or darkness.

In light-grown Col-0 cells all treatments induced significantly higher skewness of the fluorescent signal as compared to mock-treated cells. In such mock-treated cells (Fig. 5A), skewness was 0.80±0.34 (Fig. 5I; mean±SD; N=32), while after treatment with 3 µM GR24 (Fig. 5B) it was 1.73±0.23 (Fig. 5I; mean±SD; N=31), and after treatments with 25 µM GR24 (Fig. 5C) and 3 µM TIS108 (Fig. 5D) it was 1.74±0.34 (Fig. 5I; mean±SD; N=17) and 1.78±0.26 (Fig. 5I; mean±SD; N=51), respectively. It is noteworthy that the effects of all treatments were significantly different compared to the mock treatment (Fig. 5I; p=0.0000 for 3 µM GR24, 25 µM GR24, and 3 µM TIS108), though comparable to each other. The skewness of fluorescent signal from GFP-MBD lines in *max2-1* mutant background was significantly higher compared to Col-0 (Fig. 5E-H,J; p=0.0000 for all treatments), but at comparable levels within all treatments in the *max2-1* group (Fig. 5K). The increase of skewness in the Col-0 group might be relevant to the inducible increase of cortical MT anisotropy and may also underlie the intrinsically higher order of cortical MT organization of the *max2-1* mutant compared to Col-0. At the same time, it does not seem to correlate with the loosening of MT organization within the *max2-1* group after the interference with either SL signalling or biosynthesis.

**Fig. 5.**
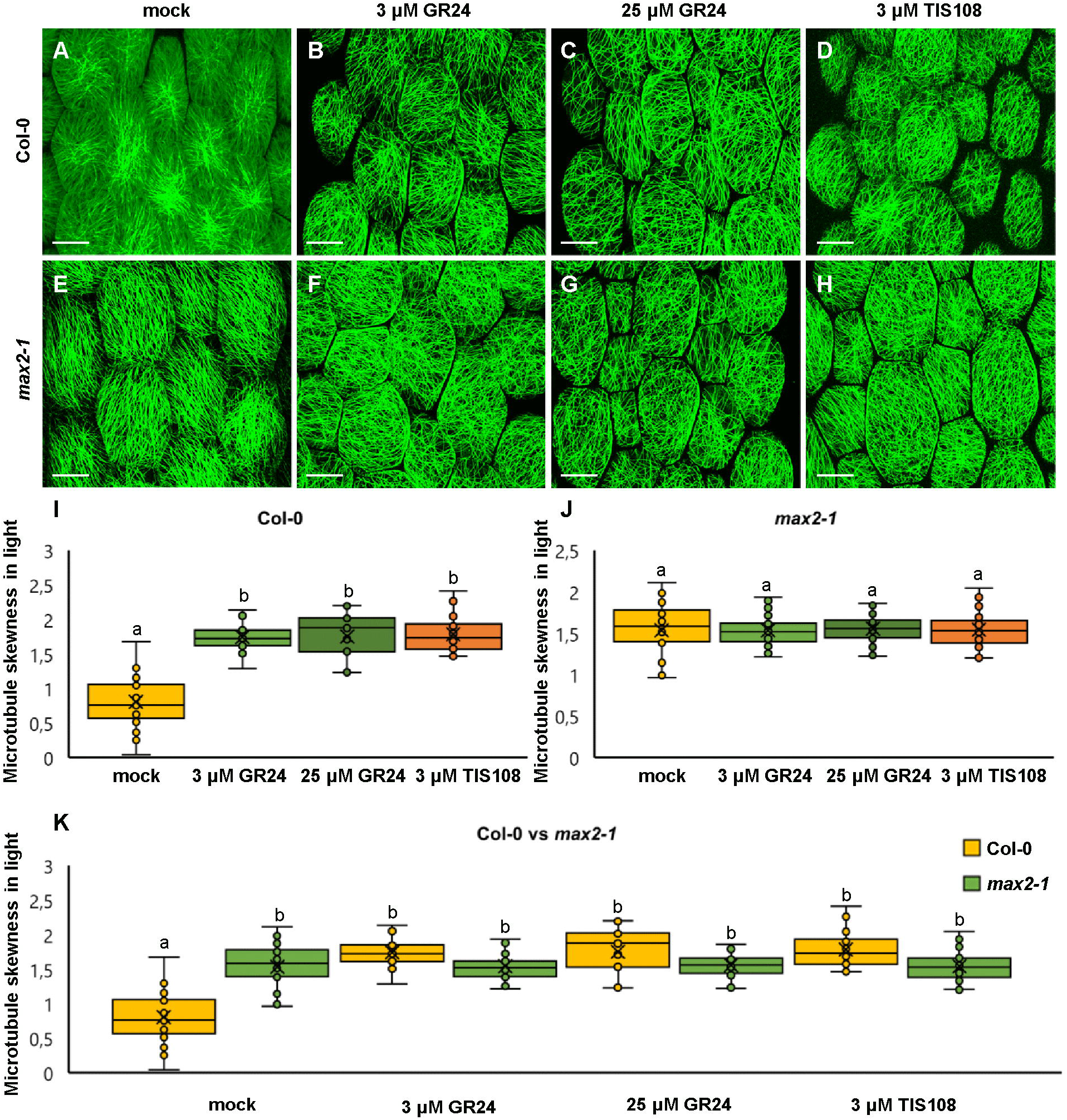
Skewness of fluorescence distribution of GFP-MBD-labelled microtubules of light- grown Arabidopsis Col-0 and *max2-1* epidermal hypocotyl cells in the presence or absence of GR24 synthetic strigolactone (3 and 25 µM) or the biosynthetic inhibitor of strigolactone production TIS108 (3 µM). (A-D) Overviews of hypocotyl of Col-0 seedlings treated with solvent alone (mock; A); 3 µM GR24 (B); 25 µM GR24 (C); 3 µM TIS108 (D). (E–H) Overview of hypocotyl of *max2-1* seedlings treated with solvent alone (mock; E); 3 µM GR24 (F); 25 µM GR24 (G); 3 µM TIS108 (H). (I, J) Quantitative assessment of fluorescence distribution skewness, comparing Col-0 (I; N≥17; Welch’s ANOVA was followed with Scheffé’s test, statistical comparison is shown within groups sharing the same genotype; letters in the graph are shared by groups without statistically significant differences at the 0.01 probability level; results are in Table S7) and *max2-1* (J; N≥34; Welch’s ANOVA showed no statistically significant difference within the dataset; F (3, 161)=0.0777, p=0.9719). (K) Collective quantification of fluorescence skewness comparing Col-0 and *max2-1* in a pairwise manner in all experimental conditions (N≥17; two-way ANOVA was followed with Scheffé’s test, statistical comparison is shown within groups sharing the same genotype; letters in the graph are shared by groups without statistically significant differences at the 0.001 probability level; results are in Table S8). In all box plots, average is presented by ^×^, median by the middle line, 1^st^ quartile by the bottom line, 3^rd^ quartile by the top line; the whiskers lie within the 1.5^×^ interquartile range (defined from the 1^st^ to the 3^rd^ quartiles) while outliers are omitted. Scale bars: 20 µm.

In dark-grown seedlings of either Col-0 or *max2-1* mutants, skewness of fluorescence distribution of GFP-MBD-labelled cortical MT was comparable between both groups with no statistically significant difference (Fig. 6). In detail, mock-treated Col-0 cells showed a skewness value of 1.49±0.3 (Fig. 6A,I; mean±SD; N=57), while 1.52±0.3 (Fig. 6B,I; mean±SD; N=58) – after the treatment with 3 µM GR24, 1.58±0.36 (Fig. 6C,I; mean±SD; N=24) – after 25 µM GR24, and 1.55±0.31 (Fig. 6D,I; mean±SD; N=22) – after 3 µM TIS108. Similarly, etiolated mock-treated *max2-1* seedlings (Fig. 6E) showed fluorescence skewness of 1.71±0.31 (Fig. 6J; mean±SD; N=31), 1.71±.30 (Fig. 6J; mean±SD; N=39) after treatment with 3 µM GR24 (Fig. 6F), 1.73±0.29 (Fig. 6J; mean±SD; N=34) after 25 µM GR24 (Fig. 6G), and 1.71±0.35 (Fig. 6J; mean±SD; N=47) after 3 µM TIS108 (Fig. 6H). Although skewness values of *max2-1* etiolated seedlings were consistently higher than those of the Col-0 group, the differences inferred were not significant (Fig. 6K). These results are partially consistent with the mild effects of exogenous SL or SL biosynthesis inhibitor on the anisotropy of MT organization in etiolated seedlings.

**Fig. 6.**
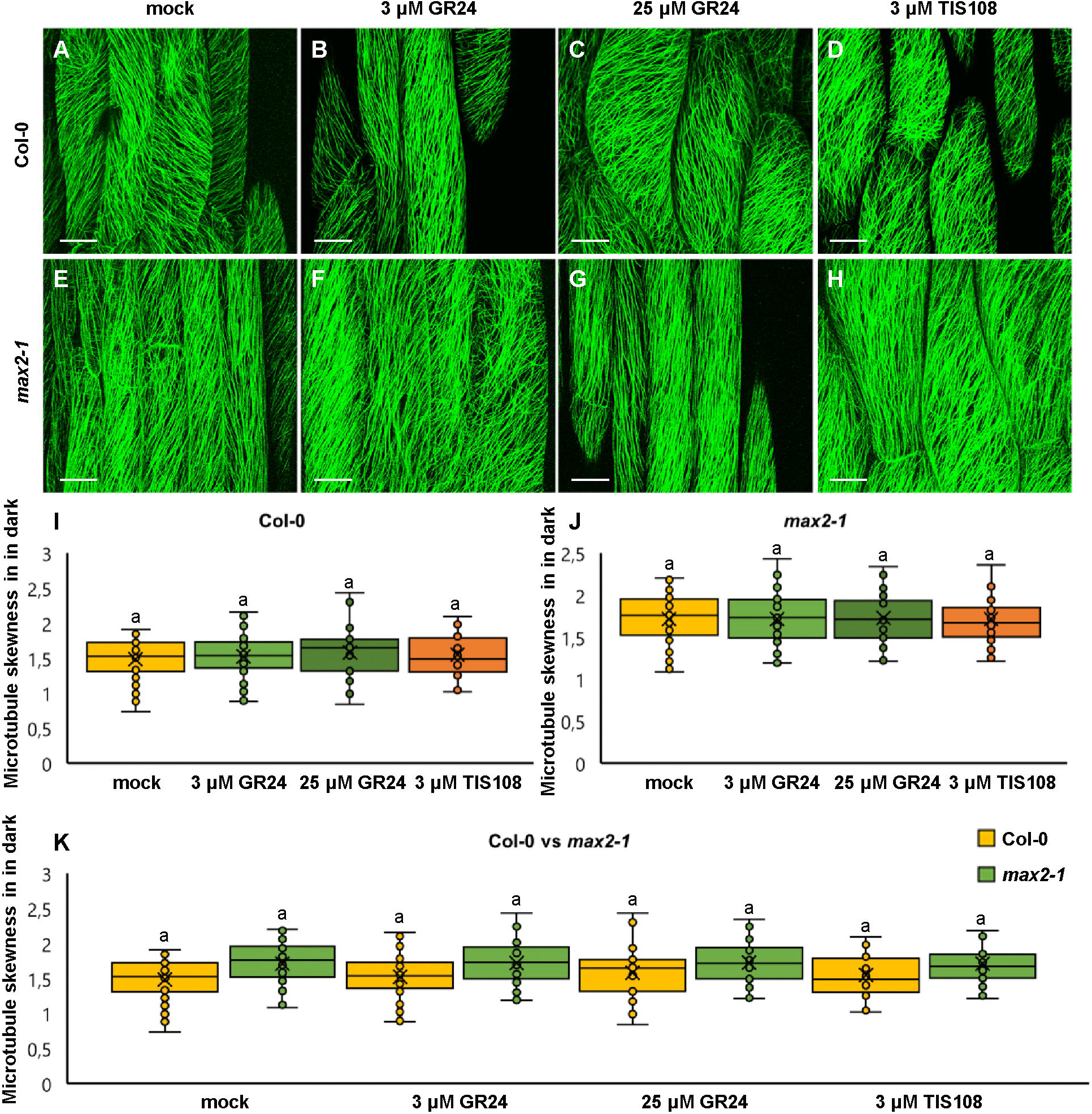
Skewness of fluorescence distribution of GFP-MBD-labelled microtubules of etiolated Arabidopsis Col-0 and *max2-1* epidermal hypocotyl cells in the presence or absence of GR24 synthetic strigolactone (3 and 25 µM) or the biosynthetic inhibitor of strigolactone production TIS108 (3 µM). (A–D) Overviews of hypocotyl of Col-0 seedlings treated with solvent alone (mock; A); 3 µM GR24 (B); 25 µM GR24 (C); 3 µM TIS108 (D). (E– H) Overview of hypocotyl of *max2-1* seedlings treated with solvent alone (mock; E); 3 µM GR24 (F); 25 µM GR24 (G); 3 µM TIS108 (H). (I, J) Quantitative assessment of fluorescence distribution skewness comparing Col-0 (I; N≥22; Welch’s ANOVA showed no statistically significant difference within the dataset; F (3, 161)=0.4564, p=0.7138) and *max2-1* (J; N≥31; Welch’s ANOVA showed no statistically significant difference within the dataset; F (3, 151)=0.0161, p=0.9972). (K) Collective quantification of fluorescence skewness comparing Col- 0 and *max2-1* in a pairwise manner in all experimental conditions (N≥22; two-way ANOVA was followed with Scheffé’s test, but there was no statistically significant difference at the 0.001 probability level; results are in Table S9). In all box plots, average is presented by ×, median by the middle line, 1st quartile by the bottom line, 3^rd^ quartile by the top line; the whiskers lie within the 1.5^×^ interquartile range (defined from the 1^st^ to the 3^rd^ quartile) while outliers are omitted. Scale bars: 20 µm.

### Strigolactones are involved in the regulation of cortical microtubule dynamics

MT dynamics were followed by means of time-lapsed SIM in hypocotyl cells of dark-grown Col-0 or *max2-1* mutants both stably expressing the GFP-MBD MT marker. Using a frame rate of ca. 0.4 frames per second (fps) it was possible to record time series of end-wise length excursions of individual or bundled MT and quantify measures of plus end dynamic instability using appropriately generated kymographs.

In mock-treated Col-0 hypocotyl epidermal cells expressing GFP-MBD (Fig. 7A,B; Supplementary Movie 1) plus end growth and shrinkage rates as well as catastrophe and rescue frequencies measured from appropriate kymographs (Fig. 7C,D) were within previously published values. Briefly, the average growth rate was 5.46±2.76 µm×min-^1^ (mean±SD; N=53 MT ends), while the average shrinkage rate was 16.48±6.25 m×min^-1^ (mean±SD; N=50 MT ends). Furthermore, catastrophe frequency was 0.0122 events×sec^-1^, while rescue frequency was 0.0512 events×sec^-1^. In both cases of GR24 treatment (3 and 25 µM) plus end MT dynamics were considerably slowed during both growth and shrinkage. At the concentration of 3 µM (Fig. 7E-H, Supplementary Movie 2) the average growth rate was 2.05±0.96 µm×min^-1^ (mean±SD; N=50 MT ends), and the average shrinkage rate was 12±8 µm×min^-1^ (mean±SD; N=50 MT ends). Catastrophe frequency was 0.0082 events×sec^-1^, while rescue frequency was 0.0332 events×sec^-1^. At 25 µM (Fig. 7I-L, Supplementary Movie 3) the average growth rate was 2.01±1.23 µm×min^-1^ (mean±SD; N=59 MT ends) and the average shrinkage rate was 6.23±5.46 µm×min^-1^ (mean±SD; N=42 MT ends). Catastrophe frequency was 0.0078 events×sec^-1^, while rescue frequency was 0.0288 events×sec^-1^. The biosynthetic inhibitor TIS108 (Fig. 7M–P, Supplementary Movie 4) strongly inhibited MT plus end dynamic parameters. In general, the average growth rate was 0.70±0.32 µm×min^-1^ (mean±SD; N=33 MT ends) and the average shrinkage rate was 4.59±5.42 µm×min^-1^ (mean±SD; N=28 MT ends). Catastrophe frequency was 0.0077 events×sec^-1^ while rescue frequency was 0.0255 events×sec^-1^.

**Fig. 7.**
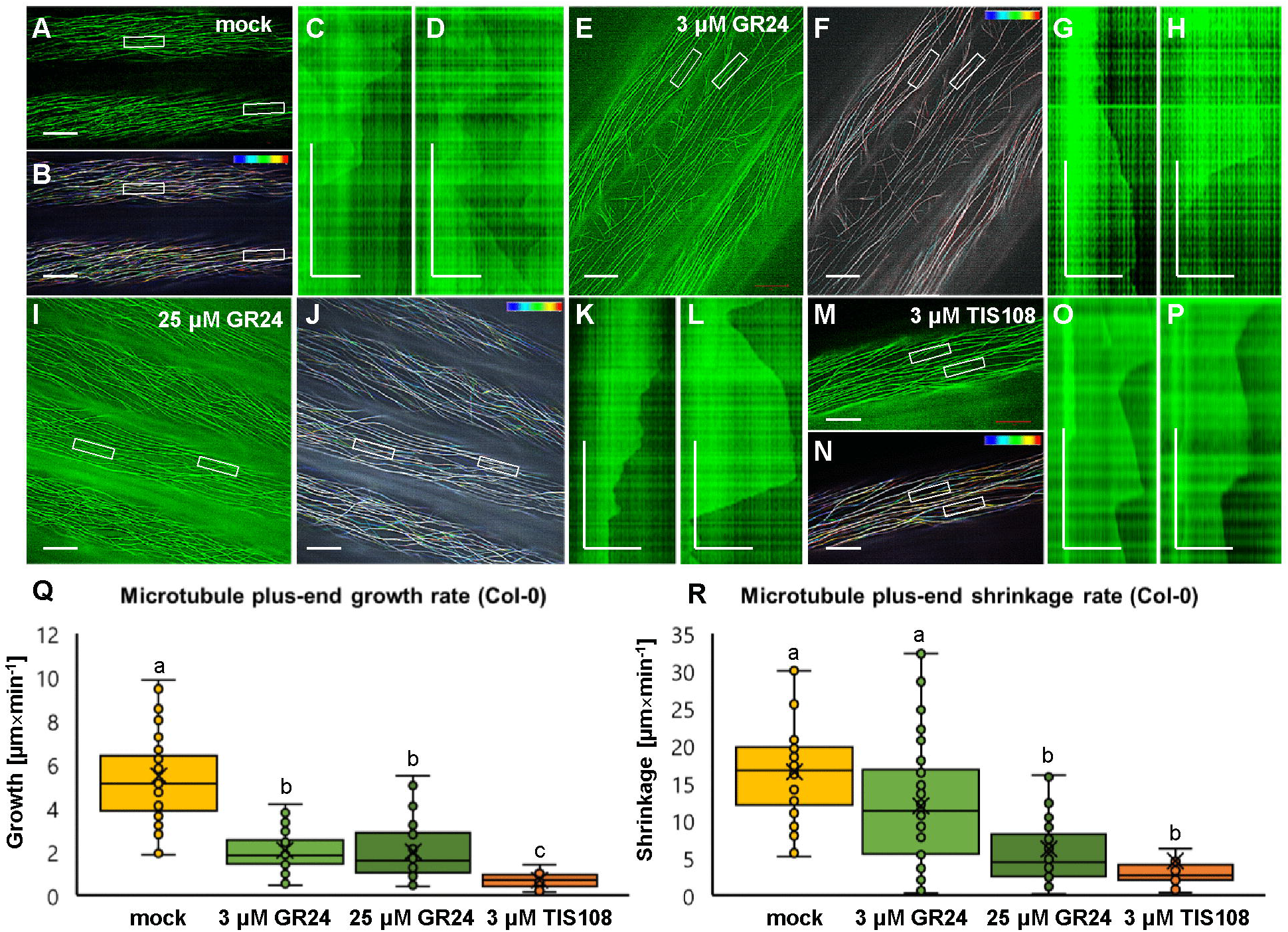
Analysis of microtubule dynamics of Arabidopsis Col-0 expressing the GFP-MBD microtubule marker in the presence or absence of GR24 synthetic strigolactone (3 µM and 25 µM) or the biosynthetic inhibitor of strigolactone production TIS108 (3 µM). (A, B) Overview (A) and color-coded projection (B) of the time series corresponding to mock-treated Col-0 (see Supplementary Movie 1). (C, D) Two kymographs showing length fluctuations of the left (C) and the right (D) boxed areas of (A, B). (E, F) Overview (E) and color-coded projection (F) of the time series corresponding to Col-0 treated with 3 µM GR24 (Supplementary Movie 2). (G, H) Two representative kymographs from boxed areas 1 and 2 of (E, F) showing decelerated and sustainable growth and shrinkage. (I, J) Overview (I) and color-coded projection (J) of the time series corresponding to Col-0 treated with 25 µM GR24 (Supplementary Movie 3). (K, L) Two representative kymographs from boxed areas 1 and 2 of (I, J) showing prolonged growth and shrinkage at lower rates compared to mock-treated cells. (M, N) Overview (M) and color- coded projection (N) of the time series corresponding to Col-0 treated with 3 µM TIS108 (Supplementary Movie 4). (O, P) Two representative kymographs from boxed areas 1 and 2 of (M, N) showing prolonged growth and shrinkage at lower rates compared to mock-treated cells. (Q, R) Quantitative assessment of microtubule growth (Q; N≥33; Welch’s ANOVA was followed with Scheffé’s test, statistical comparison is shown within groups sharing the same genotype; letters in the graph are shared by groups without statistically significant differences at the 0.01 probability level; results are in Table S10) and shrinkage (R; N≥28; Welch’s ANOVA was followed with Scheffé’s test, statistical comparison is shown within groups sharing the same genotype; letters in the graph are shared by groups without statistically significant differences at the 0.01 probability level; results are in Table S11) of Col-0 GFP-MBD labeled microtubule in all experimental conditions. In all box plots, average is presented by ^×^, median by the middle line, 1st quartile by the bottom line, 3rd quartile by the top line; the whiskers lie within the 1.5^×^ interquartile range (defined from the 1^st^ to the 3^rd^ quartile) while outliers are omitted. Scale bars: 10 µm (A, B, E, F, I, J, M, N); 5 µm (C, D, G, H, K, L, O, P). All time bars correspond to 2 min.

By comparison to mock-treated cells, both parameters of MT dynamics were in most cases significantly reduced in all treatments tested (Fig. 7Q for growth rate and Fig. 7R for shrinkage rate). In terms of growth rate (Fig. 7Q) both concentrations of GR24 showed comparable reduction as compared to mock treatment, while growth rates were even more reduced in the case of treatment with TIS108 (Fig. 7Q; p=0.0000 for 3 µM GR24; 25 µM GR24 and 3 µM TIS108). Shrinkage rates were also reduced in all treatments (Fig. 7R; p=0.0108 for 3 µM GR24; and p=0.0000 for both 25 µM GR24 and 3 µM TIS108).

The most striking feature of GFP-MBD MT in the *max2-1* mutant was the significantly lower growth rate and most importantly the long-sustained growth periods of nearly every MT examined. The prolonged elongation of cortical MT was clearly evident in mock-treated *max2-1* seedlings (Fig. 8A–E, Supplementary Movie 5) with the average growth rate being 2.09±1.27 µm×min^-1^ (mean±SD; N= 32 MT ends) and the average shrinkage rat being 8.48±7.06 µm×min^-1^ (mean±SD; N= 54 MT ends). In such seedlings, the catastrophe frequency was 0.0087 events×sec^-1^ and the rescue frequency was 0.0266 events×sec^-1^. However, the exogenous application of GR24 at either 3 or 25 µM, or the treatment with TIS108, had no effect on any parameter of MT dynamics compared to mock-treated *max2-1* cells. Briefly, in *max2-1* seedlings treated with 3 µM GR24 (Fig. 8F–K, Supplementary Movie 6), the average growth rate was 2.25±1.35 µm×min^-1^ (mean±SD; N=134 MT ends) and the average shrinkage rate was 9.03±7.38 µm×min^-1^ (mean±SD; N=87 MT ends). In turn, catastrophe frequency was 0.0071 events×sec^-1^ while rescue frequency was 0.0301 events×sec^-1^. Similar was the situation of *max2-1* seedlings treated with 25 µM (Fig. 8L-P, Supplementary Movie 7) were average growth was measured at 2.02±1.73 µm×min^-1^ (mean±SD; N=167 MT ends), and average shrinkage rate was calculated to be 9.34±6.76 µm×min^-1^ (mean±SD; N=92 MT ends). Catastrophe and rescue frequencies were 0.0081 events×sec^-1^ and 0.0264 events×sec^-1^, respectively. As in the case of GR24, *max2-1* mutants were relatively insensitive to TIS108 treatment as well (Fig. 8Q–U, Supplementary Movie 8). Therefore, the growth rate was 2.13±1.24 µm×min^-1^ (mean±SD; N=41 MT ends) and the shrinkage rate was 10.84±7.70 µm×min^-1^ (mean±SD; N=20 MT ends). Catastrophe and rescue frequencies were 0.0083 events×sec^-1^ and 0.0222 events×sec^-1^, respectively. As mentioned before, treatments had no significant effect on neither growth (Fig. 8V), nor shrinkage (Fig. 8W) within the *max2-1* group.

**Fig. 8.**
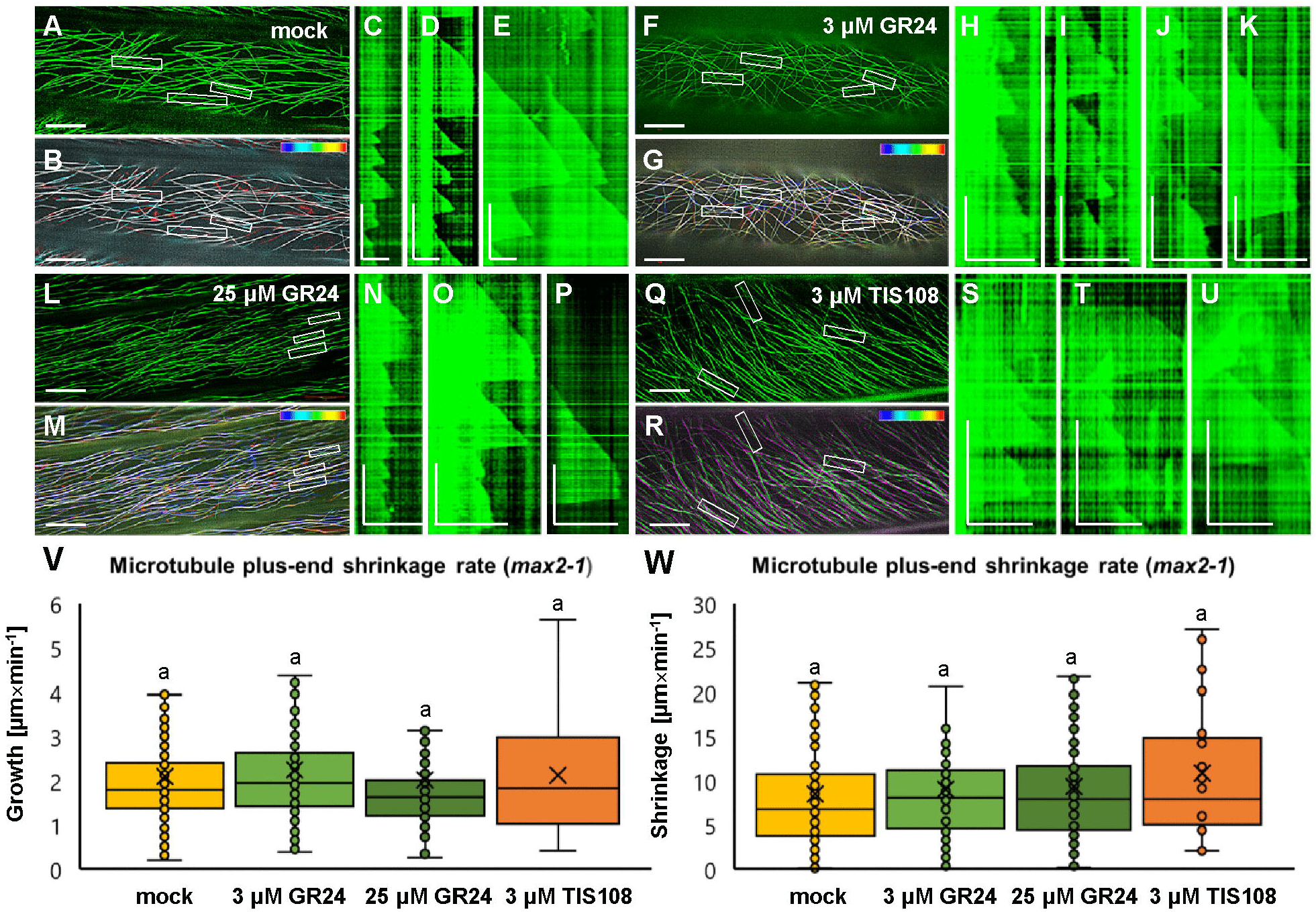
Analysis of microtubule dynamics of Arabidopsis *max2-1* mutant expressing the GFP-MBD microtubule marker in the presence or absence of GR24 synthetic strigolactone (3 and 25 µM) or the biosynthetic inhibitor of strigolactone production TIS108 (3 µM). (A, B) Overview (A) and color-coded projection (B) of the time series corresponding to mock-treated *max2-1* (Supplementary Movie 5). (C, D, E) Three kymographs showing microtubule length fluctuations corresponding to boxed areas 1,2,3 of (A, B), indicative of slower and prolonged growth and shrinkage compared to Col-0. (F, G) Overview (F) and color-coded projection (G) of the time series corresponding to *max2-1* treated with 3 µM GR24 (Supplementary Movie 6). (H–K) Four representative kymographs from boxed areas 1,2,3 and 4 of (F, G) showing similar microtubule dynamics as in mock-treated cells. (L, M) Overview (L) and color-coded projection (M) of the time series corresponding to *max2-1* treated with 25 µM GR24 (Supplementary Movie 7). (N–P) Three representative kymographs from boxed areas 1,2 and 3 of (L, M) showing comparable growth and shrinkage to mock-treated cells. (Q, R) Overview (Q) and color-coded projection (R) of the time series corresponding to *max2-1* treated with 3 µM TIS108 (Supplementary Movie 8). (S-U) Three representative kymographs from boxed areas 1,2 and 3 of (Q, R). (V,W) Quantitative assessment of microtubule growth (Q; N≥41; Welch’s ANOVA showed no statistically significant difference within the dataset; F (3, 601)=0.6081, p=0.6106) and shrinkage (R; N≥20; Welch’s ANOVA showed no statistically significant difference within the dataset; F (3, 333)=80.2659, p=0.6649) of GFP-MBD labelled microtubule in all experimental conditions. In all box plots, average is presented by ×, median by the middle line, 1st quartile by the bottom line, 3^rd^ quartile by the top line; the whiskers lie within the 1.5× interquartile range (defined from the 1^st^ to the 3^rd^ quartiles) while outliers are omitted. Scale bars: 10 µm (A, B, F, G, L, M, Q, R); 5 µm (H–K, N–P, S–U); 2 µm (C–E). All time bars correspond to 2 min.

Uniformly, growth rates in Col-0 group were reduced compared to mock treatment in a similar manner to the growth rates of *max2-1* (Supplementary Fig. 2A; p=0.0000 for 3 µM GR24, 25 µM GR24, and 3 µM TIS108). Reductions of shrinkage rates showed higher variability either comparing different experimental conditions within the Col-0 group, or by comparing the Col-0 group with the *max2-1* group (Supplementary Fig. 2B; p=0.1732 for 3 µM GR24; and p=0.0000 for both 25 µM GR24 and 3 µM TIS108).

Conclusively, the aforementioned results suggest that alterations in SL signaling either by chemical (GR24 and TIS108 treatments) or genetic (*max2-1* mutant) interference, uniformly reduce MT dynamicity and likely promote MT longevity, as evidenced by the considerably lower catastrophe frequencies observed.

## Discussion

Being produced mainly in the roots (Foo *et al*., 2013) SL adjust both shoot (Gomez-Roldan *et al*., 2008; Umehara *et al*., 2008) and root (Ruyter-Spira *et al*., 2011) development in vascular plants as well as in moss caulonema (Hoffman *et al*., 2014) to changing environmental conditions. Early grafting experiments showed that SL are transported from roots to shoot in the xylem of Arabidopsis and tomato, which provided insight into SL signalling regulation via localization and transport (Kohlen *et al*., 2011). SL may enhance and inhibit organ size and number depending upon the organ (rhizoid or caulonema; Hoffmann *et al*., 2014). In this study, the role of SL in shaping shoot architecture via microtubular cytoskeleton rearrangement was addressed. However, the detailed mechanisms governing their regulation of plant development still remain to be elucidated.

The exogenous application of a synthetic SL (GR24; Umehara *et al*., 2008) and an inhibitor of endogenous SL production (TIS108; a potent triazole-containing inhibitor of cytochrome P450 monooxygenases; Ito *et al*., 2010; 2011; 2013), resulted in hypocotyl growth alterations in both Col-0 and a SL perception mutant in *MAX2*, a gene encoding a member of the F-box leucine-rich repeat protein family which is likely the substrate recognition subunit of SCF ubiquitin E3 ligase for targeted proteolysis at the proteasome (Stirnberg *et al*., 2002; Wang Y *et al*., 2013; Wang L *et al*., 2015). Alleles of *max2* mutant are rendered insensitive to exogenous SL application in phenomena such as SL-induced inhibition of hypocotyl elongation (Jia *et al*., 2014; Wang L *et al*., 2020), suppression of shoot branching (Wang Y *et al*., 2013; Liu *et al*., 2014; Li *et al*., 2016) and lateral root formation (Ruyter-Spira *et al*., 2011; Li *et al*., 2016). Moreover, recent works associated the function of MAX2 with photomorphogenesis (Lopez-Obando *et al*., 2018).

Previous studies on the effects of exogenous SL on vegetative growth have shown that compounds such as GR24 exert an inhibitory role on the skotomorphogenic elongation of the hypocotyl and on branching processes of either the shoot or the root culminating in the reduction of tilling and lateral root formation among others (Ruyter-Spira *et al*., 2013; Jiang *et al*., 2016; Sun *et al*., 2019).

The effects of SL signalling manipulation were conspicuously evident in light-grown and, to lesser extent, in etiolated seedlings. Indeed, previous studies have shown that exogenous SL application halts hypocotyl elongation of light-grown seedlings in a dose-dependent manner, being notable at even lower concentrations as the ones used herein (e.g., at 100 nM; Jia *et al*., 2014). Importantly, *max2* mutant alleles show negligible response at low concentrations of exogenous SL and exhibited inhibition of hypocotyl elongations at concentrations exceeding 25 µM (Jia *et al*., 2014). These results corroborate the previous studies on the synergy between SL and light perception (Brewer *et al*., 2013) involving a correlation of SL perception with both phytochrome and cryptochrome light-dependent signalling (Jia *et al*., 2014).

Diffuse organ growth (i.e., elongation or lateral expansion) is conditionally regulated by physical or hormonal signals and involves the positional control of cellulose microfibril deposition. In this sense, cortical MT have been repeatedly shown to underlie cell and organ growth rate and directionality as shown in the case of light (e.g., Sambade *et al*., 2012; Lindeboom *et al*., 2013; Ma *et al*., 2018), mechanical stimulation (Louveaux *et al*., 2016; Takatani *et al*., 2020), and hormonal cues including ethylene (Ma *et al*., 2018; Wang X *et al*., 2020), auxin (True and Shaw, 2020) and gibberellins (Vineyard *et al*., 2013; Locascio *et al*., 2013).

In light of the above, the present study was extended to address whether manipulation of SL signalling could be related to cytoskeletal remodelling, thus, the organization and the dynamics of cortical MT were studied in appropriate fluorescent marker lines of both Col-0 and *max2-1* mutants. In terms of organization, exogenous SL application and inhibition of endogenous SL biosynthesis under standard light/dark exposure did not affect significantly cortical MT orientation in Col-0 but had a prominent effect in *max2-1* mutants, promoting randomization of the cortical array. By contrast, MT bundling was enhanced after all treatments in Col-0 but remained unchanged in *max2-1* mutants, which seemingly exhibited a higher level of bundling than Col-0 in all circumstances. Notably, such MT organization features as ordering and bundling remained fairly unresponsive to the chemical treatments in etiolated seedlings of both Col-0 and *max2-1*. MT dynamics were considerably lowered after chemical manipulation of SL signalling in Col-0, while the inherently lower MT dynamics of *max2-1* remained unresponsive to GR24 and TIS108.

Owing to the previous connection of SL with phytochrome and cryptochrome light perception pathways, the differential responses of cortical MT to SL content alterations under light or dark growth conditions is expected. Earlier studies have already demonstrated the interdependence between phytochromes and light-induced MT reorientation (Fischer and Schopfer, 1997), while more recently, the reorientation of cortical MT under blue light stimulation was attributed to the stimulation of KATANIN-mediated MT severing via the activation of the PHOT1 and PHOT2 phototropin photoreceptors (Lindeboom *et al*., 2013).

At present, the molecular components responsible for SL-mediated suppression of MT dynamics in Arabidopsis remain unknown. Its putative mechanisms are summarized in the hypothetical model of the interplay of light- and SL-induced pathways, which regulates the organization and dynamics of cortical microtubules resulting in the subsequent changes of hypocotyl growth and morphology (Fig. 9). The initial perception of SL in karrikin-independent pathway is provided by α/β hydrolase AtD14 (Seto et al., 2019), being activated by its binding with the ligand and able to form complex with one of the F-box protein MAX2 (reviewed by Kumar *et al*., 2015a; Wang L *et al*., 2020; Yoneyama *et al*., 2020). Upon the assembly of the SCF complex including CULLIN1 (CUL1), Skp1 (S-phase kinase-associated protein 1) and E3 ubiquitin-protein ligase RING-BOX1 (RBX1) it directs ubiquitin transfer from an E2 ligase onto target proteins, which leads to their proteasome degradation. SCF complex containing MAX2 is known to affect plant development via the degradation of SUPPRESSOR OF MORE AXILLARY GROWTH2-LIKE (SMLX) proteins (Wang L *et al*., 2020). Another putative target protein for MAX2-mediated ubiquitination is one the key transcription factors of the brassinosteroid pathway, namely BRASSINAZOLE-RESISTANT 1 (BZR1), which directly targets and upregulates MICROTUBULE DESTABILIZING PROTEIN40 (MDP40), a positive regulator of hypocotyl cell elongation by altering the stability of cortical microtubules (Wang *et al*., 2012). The more pronounced randomization of cortical MT array, increased MT bundling and stabilization as well as reduced MT dynamicity and likely promoted MT longevity leading to the stalled hypocotyl elongation and mild radial swelling of epidermal cells might be regulated by this BZR1- MDP40 pathway branch as well.

**Fig. 9.**
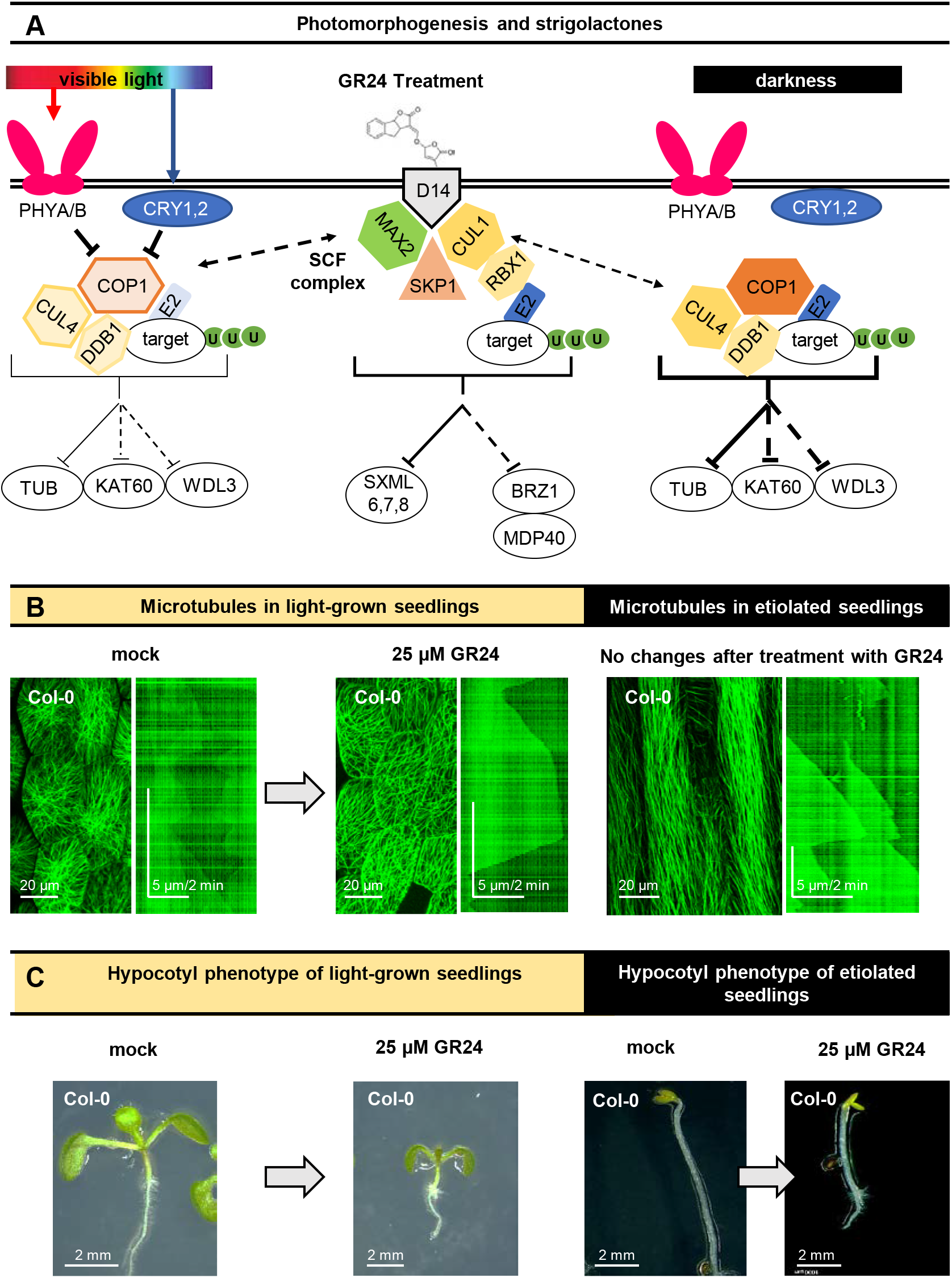
Hypothetical model of light-dependent strigolactone effects on microtubules in Arabidopsis. (A) Red and blue light is perceived by PHYTOCHROMES A and B (PHYA/B) and CRYPTOCHROMES 1 and 2 (CRY1/2), respectively, which inhibit the E3 ligase complex consisting of CONSTITUTIVE PHOTOMORPHOGENIC 1 (COP1), CULLIN4 (CUL4) and DAMAGE-BINDING PROTEIN 1 (DDB1). This E3 ligase complex directs the transfer of ubiquitin (U) from an E2 ligase onto targets, which generally leads to their degradation by proteasomes. Bellow the complex, known (solid line) and putative (dotted line) targets are shown, specifically TUBULIN (TUB), KATANIN 60 (KAT60) and WAVE-DAMPENED 2- LIKE 3 (WDL3). The function of this E3 ligase complex is more prominent under darkness, when it is not inhibited by PHYA/B and CRY1/2. It has been previously proposed that COP1 might be regulated by the SKP1-CULLIN-F-BOX (SCF) complex containing an F-box protein MORE AUXILARY GROWTH 2 (MAX2). This SCF complex consists of MAX2, hydrophobic scaffold protein CULLIN1 (CUL1), S-phase kinase-associated protein 1 (SKP1), and E3 ubiquitin-protein ligase RING-BOX1 (RBX1), it also functions as an E3 ligase, known (solid line) and putative (dotted line) targets are shown, namely SUPPRESSOR OF MORE AXILLARY GROWTH2-LIKE (SMLX) proteins and transcriptional repressor BRASSINAZOLE-RESISTANT 1 (BZR1), which is involved in regulation of microtubules via the MICROTUBULE DESTABILIZING PROTEIN40 (MDP40). SCF complex is activated by artificial strigolactones GR24+ binding to an α/β-hydrolase D14, strigolactone-specific receptor. (B) In light-grown seedlings the treatment with GR24, or an inhibitor of strigolactone biosynthesis (TIS108) leads to changes in microtubule organisation and dynamics: (1) more pronounced randomization of cortical microtubule array; (2) increased microtubule bundling and stabilization; (3) reduced microtubule dynamicity and likely promoted microtubule longevity. On the other hand, no significant microtubule changes were noted after similar treatments in etiolated seedlings as the trend is to maintain highly organized systems of parallel microtubules. (C) Regarding the overall hypocotyl phenotype, the strigolactone treatment inhibits hypocotyl growth and cause slight radial swelling of epidermal cells in light-grown seedlings; however, the dark-grown ones were more resistant to the changes of strigolactone content.

Alternatively, SL pathway might interplay with the light-induced one via the different type of an E3 ligase complex consisting of CUL4, DAMAGE-BINDING PROTEIN 1 (DDB1) and CONSTITUTIVE PHOTOMORPHOGENIC 1 (COP1). The COP1 is subjected to regulation by PHYTOCHROME A and B (PHYA/B), photoreceptors of red light, and CRYPTOCHROMES 1 and 2 (CRY1/2), photoreceptors of blue light (Podolec and Ulm, 2018). It has been previously proposed that COP1 might be regulated by the SCF complex containing MAX2 (Jia *et al*., 2014). Moreover, E3 ligase complex including COP1 might target tubulin (Khanna *et al*., 2014) as well as the proteins involved in cytoskeleton regulation such as phototropin-stimulated microtubule severing protein katanin (Lindeboom *et al*., 2013) and microtubule-associated protein WAVE-DAMPENED 2-LIKE 3 (WDL3) that binds to, bundles and stabilizes microtubules (Liu *et al*., 2013; Lian *et al*., 2017). Future studies should elucidate the relationship between these two complexes.

However, another plausible explanation may refer to the physiological differences between hypocotyls and roots, especially in relation to the interplay between SL signalling and light perception. As mentioned previously, light-induced MT reorientations in aboveground tissues have been shown to correlate with phytochrome (Zandomeni and Schopfer, 1993; Fischer and Schopfer, 1997) and phototropin (Lindeboom *et al*., 2013) signalling. The roots are also not indifferent to light, since dim light gradients may form at shallow depths of the soil and probably express specialized photoreceptors responsive to low illumination rates especially at the blue wavelength range (Galen *et al*., 2007 and references therein). Differences in photoreception between aboveground and soil-residing plant parts may explain discrepancies in the cellular responses to SL or SL inhibitors and this is a matter that deserves to be followed up.

Although TIS108 is an inhibitor of P450 cytochrome monooxygenases and thus antagonist of SL function, previous reports have confirmed its inhibitory effect to hypocotyl elongation (Kawada *et al*., 2019). On this basis, the follow-up effects of TIS108 on cortical microtubules organization and dynamics are in line with its observed effects on hypocotyl growth. Since the effects of TIS108 are also differentiated between light-grown and etiolated seedlings, it is likely that the TIS108-induced cytoskeletal remodelling is also associated to imbalances in SL signalling.

In conclusion, SL have robust effects on several aspects of plant development including size regulation and growth directionality of both the hypocotyl and the root. Moreover, SL reportedly act in concert with other environmental and intrinsic factors, exerting effects in the same aspects of plant development. In the search of cellular mechanisms underlying developmental implications of SL signalling, the present study highlights the significance of cytoskeletal remodelling in the process of SL-mediated inhibition of hypocotyl growth and reveals the differential regulation of both MT organization and dynamics by SL at different illumination regimes. It is reasonable to assume that SL signalling will diversely affect the growth of different plant parts with exposure to different light conditions.

## Abbreviations

BZR1: BRASSINAZOLE-RESISTANT 1
CLSM: confocal laser scanning confocal microscopy
COP1: CONSTITUTIVE PHOTOMORPHOGENIC 1
CRY1/2: CRYPTOCHROMES 1 and 2
CUL: CULLIN
DDB1: DAMAGE-BINDING PROTEIN 1
MAX2: MORE AXILLARY GROWTH2
MBD: MT-binding domain of mammalian non- neuronal MICROTUBULE ASSOCIATED PROTEIN4
MDP40: MICROTUBULE DESTABILIZING PROTEIN40
MS: Murashige and Skoog
MT: microtubules
PHYA/B: PHYTOCHROME A and B
RBX1: RING-BOX1
SCF complex: SKP1-CULLIN-F-BOX complex
SIM: structured illumination microscopy
Skp1: S-phase kinase-associated protein 1
SL: strigolactone(s)
SMLX: SUPPRESSOR OF MORE AXILLARY GROWTH2-LIKE
TUA6: α-TUBULIN 6
WDL3: WAVE-DAMPENED 2-LIKE 3.

## Supplementary data

Supplementary data are available at *JXB* online.

*Fig. S1*. Quantitative extent of the effect of strigolactone interference in hypocotyl elongation of light-grown Col-0 (A), etiolated Col-0 (B), light-grown *max2-1* (C), and etiolated *max2-1* (D) seedlings. *, p<0.05; ***, p<0.001 according to Student’s t-test.

*Fig. S2*. Pairwise comparison of Col-0 and *max2-1* microtubule plus end growth (A; (N≥33; two-way ANOVA was followed with Scheffé’s test, statistical comparison is shown within groups sharing the same genotype; letters in the graph are shared by groups without statistically significant differences at the 0.001 probability level; results are in Table S12) and shrinkage (B; N≥20; two-way ANOVA was followed with Scheffé’s test, statistical comparison is shown within groups sharing the same genotype; letters in the graph are shared by groups without statistically significant differences at the 0.001 probability level; results are in Table S13) under all experimental conditions used herein. In all box plots, average is presented by ×, median by the middle line, 1st quartile by the bottom line, 3^rd^ quartile by the top line; the whiskers lie within the 1.5^×^ interquartile range (defined from the 1^st^ to the 3^rd^ quartile) while outliers are omitted.

*Supplementary Movie S1*. SIM time series corresponding to Fig. 7A. Microtubule dynamics of etiolated, mock-treated Col-0 hypocotyl epidermal cells expressing the GFP-MBD marker.

*Supplementary Movie S2*. SIM time series corresponding to Fig. 7E. Microtubule dynamics of etiolated Col-0 hypocotyl epidermal cells expressing the GFP-MBD marker treated with 3 µM GR24.

*Supplementary Movie S3*. SIM time series corresponding to Fig. 7I. Microtubule dynamics of etiolated Col-0 hypocotyl epidermal cells expressing the GFP-MBD marker treated with 25 µM GR24.

*Supplementary Movie S4*. SIM time series corresponding to Fig. 7M. Microtubule dynamics of etiolated Col-0 hypocotyl epidermal cells expressing the GFP-MBD marker treated with 3 µM TIS108.

*Supplementary Movie S5*. SIM time series corresponding to Fig. 8A. Microtubule dynamics of etiolated, mock-treated max2-1 hypocotyl epidermal cells expressing the GFP-MBD marker.

*Supplementary Movie S6*. SIM time series corresponding to Fig. 8F. Microtubule dynamics of etiolated *max2-1* hypocotyl epidermal cells expressing the GFP-MBD marker treated with 3 µM GR24.

*Supplementary Movie S7*. SIM time series corresponding to Fig. 8L. Microtubule dynamics of etiolated *max2-1* hypocotyl epidermal cells expressing the GFP-MBD marker treated with 25 µM GR24.

*Supplementary Movie S8*. SIM time series corresponding to Fig. 8Q. Microtubule dynamics of etiolated *max2-1* hypocotyl epidermal cells expressing the GFP-MBD marker treated with 3 µM TIS108.

*Table S1*. Statistical analysis for Fig. 1Q.

*Table S2*. Statistical analysis for Fig. 1R.

*Table S3*. Statistical analysis for Fig. 2I.

*Table S4*. Statistical analysis for Fig. 3Q.

*Table S5*. Statistical analysis for Fig. 3R.

*Table S6*. Statistical analysis for Fig. 4R.

*Table S7*. Statistical analysis for Fig. 5I.

*Table S8*. Statistical analysis for Fig. 5K.

*Table S9*. Statistical analysis for Fig. 6K.

*Table S10*. Statistical analysis for Fig. 7Q.

*Table S11*. Statistical analysis for Fig. 7R.

*Table S12*. Statistical analysis for Fig. S2A.

*Table S13*. Statistical analysis for Fig. S2B.

## Acknowledgments

We gratefully acknowledge the gift of *max2-1* mutant by Prof. Hinanit Koltai (Institute of Plant Sciences ARO, Volcani Center, Bet-Dagan). We appreciate the help of Pavlína Floková in the reconstruction of some of the time series acquired by SIM. The research was supported from ERDF project “Plants as a tool for sustainable global development” (No. CZ.02.1.01/0.0/0.0/16_019/0000827), and partially by the Višegrad Out-Going Scholarship for the Eastern Partnership for Post-Masters (independent research) for the academic year 2015– 2016 (ID: 51500570).

## Author contributions

YK and JŠ designed experiments with contribution from GK. YK and SH carried out all image acquisitions with the help of GK and MO. YK, SH and GK carried out all post-acquisition image processing. YK and SH carried out all hypocotyl length and width measurements. GK acquired all necessary measurements and analyzed all data related to microtubule organization and dynamics. TV carried out statistical analyses. TP synthesized GR24. YK and GK drafted the manuscript. GK compiled all figures with input from YK, TV, and JŠ. JŠ provided funding and infrastructure.

## Data Availability Statement

All data supporting the findings of this study are available within the paper and within its supplementary materials published online.

